# A novel mouse model recapitulates features of the neurodevelopmental disorder 3p25.3 microdeletion syndrome

**DOI:** 10.64898/2026.11.10.765518

**Authors:** Sophie TB Morgan, Andrew Kueh, Andrew Keniry, Samantha Eccles, Marnie E Blewitt, Hannah K Vanyai

**Affiliations:** The Walter and Eliza Hall Institute of Medical Research, Parkville, VIC, Australia; Department of Medical Biology, the University of Melbourne, Parkville, VIC, Australia; Olivia Newton-John Cancer Research Institute, Heidelberg, Melbourne, Australia; School of Cancer Medicine, La Trobe University, Bundoora, Melbourne, Australia

## Abstract

Neurodevelopmental disorders globally affect approximately 15% of children and adolescents, but many disorders do not have mouse models, thus limiting our ability to further understand them and ultimately develop treatments. In our study, we developed a novel mouse line to model 3p25.3 microdeletion syndrome (*6qE3^del/+^*) that reliably recapitulates key features of the syndrome. *6qE3^del/+^* mice display congenital heart defects, craniofacial defects, skeletal anomalies and brain changes. We also generated another neurodevelopmental disorder model for a gene often included in the deleted region, *Brpf1*, and found that congenital heart defects and craniofacial defects may be partly attributable to the heterozygous loss of *Brpf1* in *6qE3^del/+^* mice. Together our work offers new neurodevelopment models, and provides evidence that treatments developed for *Brpf1* haploinsufficiency may also benefit 3p25.3 microdeletion syndrome.

## Introduction

Neurodevelopmental disorders (NDDs) are caused by abnormal brain development, which can result in a broad range of symptoms such as cognitive, social and neurological deficits, autism spectrum disorder, attention deficit disorders, anxiety, epilepsy, distinctive facial features, congenital heart defects, hypotonia and skeletal anomalies [1]. Specific NDDs are individually rare but cumulatively are estimated to affect 15% of children and adolescents worldwide [2]. However, owing to their low individual incidence, many NDDs have not been modelled, making the development of targeted treatments unattainable without understanding the underlying basis of these individual disorders.

3p25.3 microdeletion syndrome is a rare autosomal dominant NDD that occurs when there is a heterozygous deletion within the 3p25.3 locus, with approximately 57 published cases to date [3–16]. The size and coordinates of the deletion in patients is highly variable and the deleted region can extend beyond the 3p25.3 locus, indicating that it is not the presence of specific repeat regions that drive the loss of the locus as in the case in some other deletion/duplication syndromes [17]. The 3p25.3 locus contains several genes that have their own associated haploinsufficient NDDs and likely contribute to the clinical presentation of patients. One of these genes is the multivalent chromatin reader *BRPF1* (Bromodomain and PHD finger-containing protein 1), which is frequently included within the deleted region (24/48 of cases on ClinVar, Supplementary Figure 1A).

BRPF1 is a member of the histone acetyltransferase complex that contains either of the catalytic proteins KAT6A or KAT6B (lysine (K) acetyltransferase 6A or 6B). We have previously modelled NDDs associated with heterozygous variants in KAT6A and KAT6B [18–20], which lead to Arboleda-Tham syndrome (ARTHS) and Say-Barber-Biesecker-Young-Simpson Syndrome (SBBYSS), respectively [21, 22]. *Kat6a^-/+^* and *Kat6b^-/+^*mice recapitulate phenotypes seen in their relevant human disorder, such as impaired learning and memory formation, and social deficits [18, 20]. Since BRPF1 complexes with KAT6A/B, we wanted to extend upon our prior work by investigating NDDs with *BRPF1* haploinsufficiency.

In addition to its loss in half of 3p25.3 microdeletion syndrome cases, heterozygous variants in *BRPF1* cause Intellectual Developmental Disorder with Dysmorphic Facies and Ptosis (IDDDFP). Approximately 53 cases of IDDDFP have been published since the syndrome was first reported in 2017 [23–33]. Pathogenic variants occur primarily *de novo*, and they are predicted to be loss of function variants [31]. IDDDFP and 3p25.3 microdeletion syndrome exhibit highly overlapping features including intellectual disability, global developmental delay and distinctive facial features such as ptosis and blepharophimosis [29]. Other features can include congenital heart defects, skeletal/muscular abnormalities, autism spectrum disorder, attention deficit disorders, anxiety and epilepsy [29].

Studies in mice show that BRPF1 plays important roles in embryonic development, neurodevelopment, synaptic transmission, haematopoiesis and skeletal patterning [34–38]. Complete loss of *Brpf1* is embryonic lethal in mice at E9.5-E11.5, with E9.5 embryos exhibiting abnormal neural tube closure and vascular defects in the placenta, yolk sac and embryo [35]. Additionally, these embryos have a wide range of neurological defects, such as decreased proliferation in neural progenitors [34], abnormal formation of the brain [39] and, in forebrain-specific knockout models, hypoplasia of the dentate gyrus [40]. Furthermore, behavioural testing in forebrain-specific (*Emx1-Cre* or *CaMKIIa-Cre*) heterozygous mice show that *Brpf1* haploinsufficient mice exhibit memory and learning deficits, echoing the intellectual disability seen in the human disorder [41–43].

We created a novel mouse model carrying a microdeletion at *6qE3*, syntenic to human 3p25.3, and characterised the developmental phenotypes in these mice to establish a mouse model of 3p25.3 microdeletion syndrome. Furthermore, we created a *Brpf1* heterozygous deletion strain and compared the phenotypes between the two to delineate the contribution of *Brpf1* in 3p25.3 microdeletion syndrome. 3p25.3 microdeletion syndrome mice perished shortly after birth with severe ventricular septal defects, increased brain volume and displayed skeletal and craniofacial abnormalities that are consistent with the human disorder. *Brpf1^-/+^* mice had a low incidence of ventricular septal defects and subtle craniofacial abnormalities, implying that *Brpf1* is contributing to a subset of phenotypes, but not wholly responsible for the phenotypes seen in 3p25.3 microdeletion syndrome.

## Materials and Methods

### Ethics

All animal experiments were conducted in accordance with the Australian Code for the Care and Use of Animals for Scientific Purposes, with the approval of the Walter and Eliza Hall Animal Ethics Committee (2023.045 and 2023.033).

### Generation of mice models

Two mouse models were generated by the Melbourne Advanced Genome Editing Centre (MAGEC) by injecting fertilized single-cell-stage oocytes with Cas9 mRNA and two guide RNAs as previously described [44].

To generate *C57BL/6*-*6qE3^del/+^* mice, approximately 406-kb region was deleted by designing guides that flanked Thumpd3 to Jagn1, encompassing 14 genes in total (Table 1, Figure 1). Given *C57BL/6*-*6qE3^del/+^* mice were perinatally lethal, N1 embryos were used in these experiments, all of which came from two founder males. To obtain *CB6*-*6qE3^del/+^* mice, *C57BL/6*-*6qE3^del/+^*mice were outcrossed with *BALB/c-6qE3^+^*^/+^ then backcrossed to *BALB/c-6qE3^+^*^/+^. F1, N2 and N3 animals were used in experiments.

**Figure 1.**
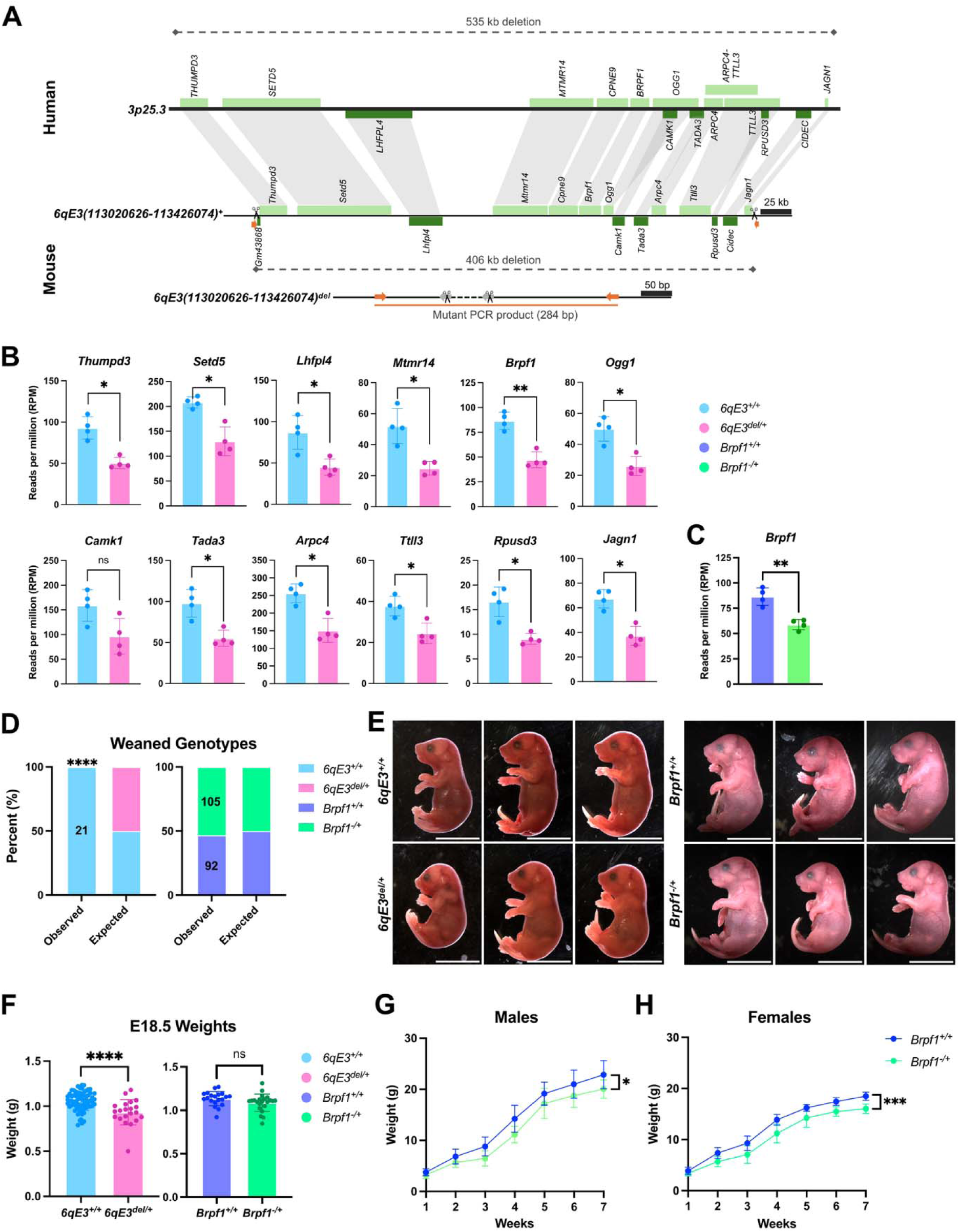
**(A)** Schematic representation of human 3p25.3 microdeletion syndrome recapitulated in mice. The 535 kb deletion in humans is equivalent to 406 kb in mice, encompassing the same 14 genes. Boxes represent genes (light green genes are transcribed on the forward strand, dark green genes transcribed on the negative strand). Scissors indicate the sgRNA cut sites, orange arrows indicate the forward and reverse primers. The *6qE3(113020626-113426074)^del^*allele (abbreviated to *6qE3^del^*) scale differs from the other schematics to illustrate the primers used for genotyping, which are upstream of each cut site (deletion region represented by dashed line, not to scale for viewing purposes). **(B-C)** RNA-Seq from **(B)** *6qE3^del/+^*and control E14.5 neural stem cells (NSCs) showing expression of genes encompassed in the deletion interval (multiple t-tests with Holm-Šidák correction, n = 2 per genotype per sex, p < 0.05*, p < 0.005**) and **(C)** expression of *Brpf1* in *Brpf1^-/+^* and control E14.5 NSCs (unpaired t-test, n = 2 per genotype per sex, p = 0.0030). **(D)** Genotype distribution of weaned pups, with observed vs. expected percentages. All *6qE3* animals that survived to weaning were wildtype (binomial test, N = 21, p < 0.0001). *Brpf1* animals were present in expected Mendelian ratio (χ*^2^* = 0.8579, df = 1, *Brpf1^+/+^*n = 92, *Brpf1^-/+^* n = 105, p = 0.3543). **(E)** Images of representative E18.5 fetuses for each genotype. Scale bar = 1 cm. **(F)** Weight of E18.5 fetuses according to genotype with ± 1 SD for *6qE3* (unpaired t-test two-tailed, *6qE3^+/+^* n = 72, *6qE3^del/+^* n = 21, p < 0.0001) and *Brpf1* (unpaired t-test two-tailed, *Brpf1^+/+^*n = 20, *Brpf1^-/+^* n = 22, p = 0.1003). Note that the *6qE3^del/+^*fetus that weighed 0.5 g was severely runted but appeared viable at time of dissection. **(G-H)** Growth curves for *Brpf1* **(G)** males and **(H) fe**males from 1-7 weeks of age. Two-way repeated measures ANOVA for genotype in **(G)** males (*Brpf1^+/+^* n = 9, *Brpf1^-/+^* n = 10, p = 0.0169) and **(H)** females (*Brpf1^+/+^* n = 8, *Brpf1^-/+^*n = 15, p = 0.0003). Error bars represent ± 1 SD.

**Table 1.**
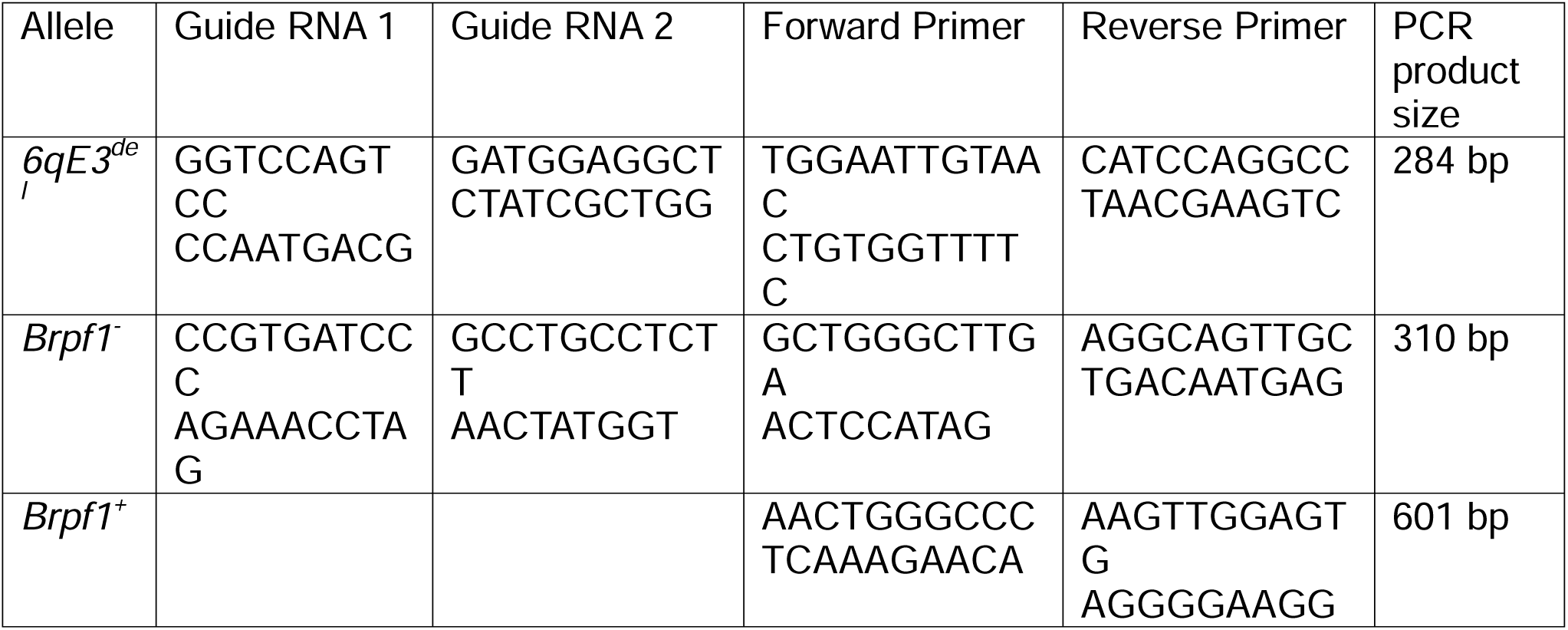
Sequences of guide RNAs and genotyping primers. All sequences are listed from 5’ to 3’.

*C57BL/6*-*Brpf1^-/+^* mice were generated by designing guide RNAs that flanked exons 4-6 to create an out-of-frame splice between exons 3 and 7, and create a loss of function allele (Table 1, Supplementary Figure 1).

### Housing

Mice were housed in ventilated cages (Tecniplast) and provided with ad libtum access to irradiated Barastoc WEHI Mouse Breeder Cubes feed (102119-1070, Ridley) and water (Hydropac, Avidity Science). Mice were kept on a 14 hour light/10 hour dark cycle.

### Plug mating

The day a vaginal plug was observed was considered embryonic day 0.5 (E0.5). Embryos were collected at ages ranging from E7.5-E18.5.

### Caesarean

Heavily pregnant mothers (E18.5-E19.5) were euthanised via cervical dislocation and litter was retrieved, cauterising the umbilical cord and clearing airways of fluid before placing on a heat pad. Pups were observed for up to 60 minutes after birth, with colour changes, establishment of breathing and survival being documented. Animals suffering respiratory distress or other illnesses were euthanised as a humane endpoint via decapitation.

### Skeletal Staining

Skeletal stains were prepared as previously described [45]. Briefly, fetuses (E17.5 or E18.5) were dissected, skinned and internal organs were removed, then briefly stored in 70-80% ethanol prior to processing. Samples were fixed for 24 hours 95% ethanol, then incubated for 24 hours in 100% acetone. Cartilage was stained using 0.03% Alcian blue (A5268, Sigma-Aldrich) in 80% ethanol and 20% glacial acetic acid, followed by two washes with 70% ethanol. Tissue was cleared using 1% KOH (w/v, in MQ-H2O) for 1 hour, then ossified bone was stained using 0.005% Alizarin red (A5533, Sigma-Aldrich) in 1% KOH. Samples were incubated in 50% glycerol:1% KOH until sufficiently cleared, then imaged in 100% glycerol.

### Histology

#### Nissl brain sections

E18.5 heads were fixed in Bouin’s fixative (HT10132-1L, Sigma-Aldrich) for 2-3 days at room temperature, washed several times with 70% ethanol, then brains were dissected, paraffin-embedded and serially sectioned into 5 um slices, then stained with cresyl violet.

#### Hematoxylin and eosion (H&E) staining

E18.5-E19.5 hearts were fixed in 4% paraformaldehyde (PFA; 158127, Sigma-Aldrich) overnight at 4 °C, stored briefly in 70-80% ethanol, rehydrated in PBS, embedded in 2% Low Melting Point agarose (LMP; ProBiogen Biochemicals), serially sectioned into 5 um slices then H&E stained. For serial head sections, E18.5 heads were fixed in 10% neutral buffered formalin (NBF) at 4 °C overnight, then stored briefly in 70% ethanol prior to processing. Heads were paraffin-embedded and sectioned into 5 um slices then H&E stained according to routine histology approaches.

### Micro-computed tomography (micro-CT)

Adult hearts were fixed in 10% NBF for 2-3 days at 4 °C. E18.5 heads were fixed in Bouin’s fixative for 2-3 days at room temperature. E18.5 hearts were fixed in 4% PFA overnight at 4 °C. Samples were stored in 70% ethanol, then rehydrated in a decreasing ethanol series before staining with 1% Lugol solution (32922, Sigma-Aldrich). Hearts were stained for 3-5 days at 4 °C, and heads were stained for 24 hours at room temperature.

After staining, all samples were embedded in 2% LMP agarose. The 3D raw data was acquired via Bruker Skyscan 1276. Each scanned sample was acquired with the X-ray source at 60 kV and the source voltage at 200 uA with a 0.5 mm aluminum filter. Each sample was rotated 180°C, with image acquisition every 0.2°C at a resolution of 10 voxels μm^3^. For volumetric measurements, the Medical Imaging Interaction Toolkit (MITK) with the AI segmentation plugin nnInteractive was used [46].Otherwise, images were visualised using Imaris.

### Immunofluorescence microscopy

Brains were fixed in 4% PFA for 1-2 days at 4 °C, then incubated with a series of solutions: 15% sucrose (w/v in PBS), 30% sucrose, 50:50 30% sucrose and Tissue-Tek® O.C.T. Compound (4583, Sakura) and 100% Tissue-Tek® O.C.T. Compound for 24 hours each at 4 °C. Brains were rapidly cryoembedded using a flash chiller (PrestoCHILL, Milestone), then cryosectioned into 16-30 μm slices using the Cryostar NX70 (Thermofisher). Slides were stored at -80 °C until ready for staining.

Slides were acclimatised to room temperature for 20 minutes, then washed thrice in PBST (0.1% Tween in PBS) before blocking and permeabilisation in 0.3% Triton X-100 (T2984, Sigma-Aldrich) and 5% Bovine Serum Albumin (BSA; 15260-037, Gibco) in PBS for 1 hour at room temperature. Samples were incubated overnight with primary antibodies in 1% BSA at 4 °C, washed thrice in PBST, stained with secondary antibodies in 1% BSA for 1 hour at room temperature protected from light, washed thrice in PBST, then stained with 0.001 mg/ml DAPI (D9542, Sigma-Aldrich) for 5 minutes. Slides were mounted using VECTASHIELD Vibrance® Antifade Mounting Medium (H1700, Vector).

All immunofluorescence samples were imaged using a Zeiss LSM 980 confocal microscope. Images were 16-bit and quantified using QuPath (v0.6.0) software. Cell segmentation was performed using a deep-learning StarDist model (dsb2018_heavy) applied to a combined, mean-transformed channel representing both DAPI and H3K9ac or H3K23ac signals. Post-processing further filtered cells by area (30-150 µm^2^), circularity (> 0.85) and DAPI parameters (mean intensity > 3000, standard deviation > 750). As such, boundaries were constrained strictly to nuclear shapes, and downstream marker quantification was extracted as the unbiased mean intensity per segmented object.

### Antibody list

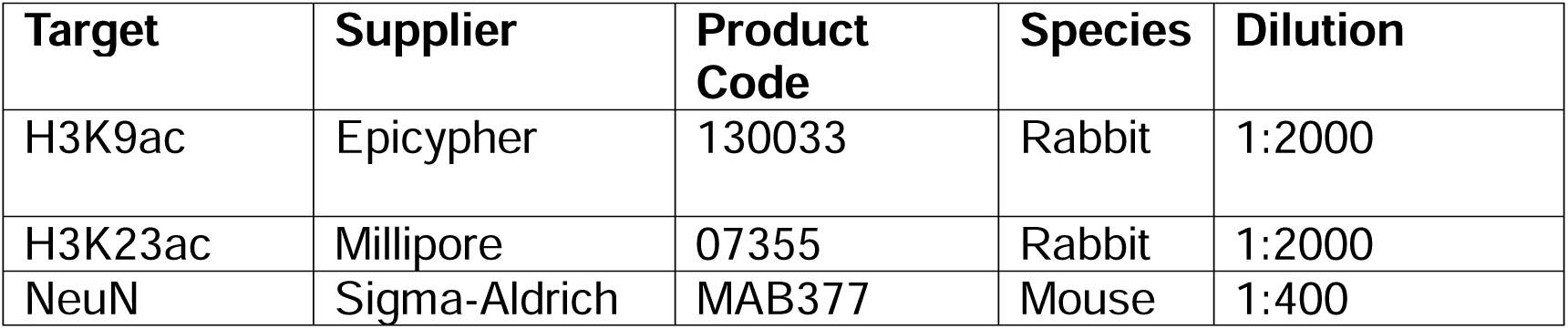

| **Target** | **Supplier** | **Product Code** | **Species** | **Dilution** |
|---|---|---|---|---|
| H3K9ac | Epicypher | 130033 | Rabbit | 1:2000 |
| H3K23ac | Millipore | 07355 | Rabbit | 1:2000 |
| NeuN | Sigma-Aldrich | MAB377 | Mouse | 1:400 |

### Neural Stem Cell Generation and Cell Culture

Neural stem cells (NSCs) were derived from the cortical regions of E14.5 mice. Tissue was cut into small pieces, then digested with 0.1 mg/ml DNase I (DN25-10MG, Sigma-Aldrich) and 0.05% trypsin-EDTA (15400-054, Gibco) in Neurocult Basal Medium (05700, StemCell Technologies) for 15-20 minutes at 37 °C. After washing thrice with media, cells were mechanically dissociated and plated in NSC complete media comprising of Neurocult Basal Medium, 5% NeuroCult™ Proliferation Supplement (05701, StemCell Technologies), 20 ng/ul EGF (02633, StemCell Technologies) and 20 ng/ul FGF (02634, StemCell Technologies) containing 5 μg/ml laminin (23017015, Life Technologies).

Cells were incubated at 37 °C with 5% CO_2_. When passaging, cells were detached using StemPro™ Accutase™ Cell Dissociation Reagent (A1110501, Gibco), quenched with media, centrifuged at 500 rpm for 5 minutes, then replated as required. NSCs were not used beyond passage 6.

### RNA Sequencing

RNA-seq was performed on RNA extracted from E14.5 mouse NSCs at passages 5-6 using the Quick-RNA Miniprep Kit (R1054, Zymo Research). Sequencing libraries were generated using the TruSeq RNA Sample Preparation Kit (Illumina) and sequenced in-house on a NextSeq 2000 platform (Illumina), producing 66 bp paired-end reads. Sequence quality assessment and adapter trimming were carried out using FastQC (https://www.bioinformatics.babraham.ac.uk/projects/fastqc/) and Trim Galore (https://github.com/FelixKrueger/TrimGalore), respectively. Reads were aligned to the mm10 mouse reference genome using HISAT2 [47]. Gene expression levels were quantified as log2RPM values using the RNA-seq quantitation pipeline in SeqMonk (https://www.bioinformatics.babraham.ac.uk/projects/seqmonk/). Downstream data exploration and visualisation were conducted in SeqMonk, and differential gene expression analysis was performed using EdgeR [48] within the SeqMonk environment. The sequencing data is accessible via GEO under the accession number GSE348152.

## Results

### Creation of a mouse model for 3p25.3 microdeletion syndrome

Mouse models are invaluable tools for studying the pathogenic mechanisms underlying human disorders and ultimately provide a whole organism system in which to test potential treatments. To develop models of the phenotypically overlapping NDDs, we created two new lines of mice, one to model 3p25.3 microdeletion syndrome and the other to model *BRPF1* haploinsufficiency.

The 3p25.3 locus is evolutionarily conserved between humans and mice, demonstrating strict gene synteny and collinearity. Given the area and size of the deletion varies significantly between patients, we decided to design a model that was wholly contained within the 3p25.3 locus that spanned over a common region of overlap and contained some of the smallest deletions reported in patients (Figure 1A).

The 3p25.3 microdeletion syndrome line (denoted *6qE3* as the equivalent locus in mice) had a heterozygous 406 kb deletion encompassing 14 genes including *Brpf1* (Figure 1A). To validate that our microdeletion recapitulates the haploinsufficiency of the disorder, we created neural stem cells (NSCs) from E14.5 fetuses to measure the expression of these genes in a defined neural cell type by RNA sequencing. Of the genes within the deletion interval that were detectably expressed, *6qE3(113020626-113426074)^del/+^*(abbreviated hereafter to *6qE3^del/+^*) NSCs showed an approximately 50% reduction in expression, with the exception of the variably expressed *Camk1* (calcium-calmodulin-dependent protein kinase I) (Figure 1B). Of genes not contained within the deleted region, *6qE3^del/+^* NSCs had very few genome-wide significant changes (Supplementary Figure 1B, Supplementary Table 1), with 50 differentially expressed (DE) genes (17 downregulated, 33 upregulated, FDR < 0.1).

*Brpf1^-/+^* mice had a heterozygous deletion of exons 4-6 (Supplementary Figure 1D). *Brpf1^-/+^* NSCs confirmed *Brpf1* downregulation (Figure 1C). Similar to *6qE3^del/+^*, *Brpf1^-/+^* NSCs had very few genome-wide significant changes, with 24 DE genes (6 downregulated, 18 upregulated, FDR < 0.1) (Supplementary Figure 1C, Supplementary Table 2). We asked if there was any overlap in the genome wide DE genes and found that two genes, *Ecscr* (endothelial cell surface chemotaxis receptor) and *Sstr3* (somatostatin receptor 3), are both upregulated in *6qE3^del/+^* and *Brpf1^-/+^* NSCs compared to wildtype (WT) NSCs which was statistically significant (hypergeometric test for overlap enrichment, p = 0.00079, 13,739 detectably expressed genes). This demonstrates that there is detectable, though subtle, overlap between the disorders at the transcriptomic level in E14.5 NSCs.

*6qE3^del/+^* animals were present at E18.5 but not at weaning, indicating lethality occurred within this period (Figure 1D). *6qE3^del/+^* E18.5 fetuses appeared smaller and weighed less than *6qE3^+/+^* (Figure 1F). We theorize that the two viable founder mice (used to generate embryos shown throughout text) exhibited mosaicism, thus enabling founder survival.

*Brpf1^-/+^* animals were present in the expected Mendelian ratio at weaning (Figure 1D). We confirmed previous reports [35] that homozygous loss of *Brpf1* is mid-gestation embryonic lethal (Supplementary Figure 1E-G). *Brpf1^-/+^* E18.5 fetuses were similar in weight to their wildtype counterparts (Figure 1F), but *Brpf1^-/+^*mice consistently weighed less than *Brpf1^+/+^* over the first 7 weeks of life (Figure 1G-H).

We successfully generated mouse models for 3p25.3 microdeletion syndrome and IDDDFP, where we transcriptomically validated the heterozygous loss of our target genes. Given that we did not identify *6qE3^del/+^*animals in our weaned postnatal population, we focused on the embryonic period to further characterise the model.

### *6qE3^del/+^* pups have severe congenital heart defects consistent with 3p25.3 microdeletion syndrome human patients

The absence of *6qE3^del/+^* animals in the weaned population despite being present at E18.5 suggested lethality of the animals was occurring between late gestation and weaning. Neonatal lethality is a common phenotype in mutant mice as they must adjust from the *in utero* environment to life outside the womb [49]. In particular, the switch from placental-to-lung based oxygenation of blood is a major hurdle at this stage. Therefore, we performed caesareans on E18.5-E19.5 litters. This allowed us to see if *6qE3^del/+^* pups attempted to and established regular breathing and blood oxygenation, as indicated by pup colour transitioning from blue to pink. While all pups delivered alive made an observable attempt to breathe, only 17% of *6qE3^del/+^* pups compared to 85% of WT survived until the end of the 60 minute observation period, with majority of the *6qE3^del/+^*pups dying in the first 20 minutes (Figure 2A). 25% of the *6qE3^del/+^* pups were stillborn. An enriched fraction of live mutants compared to wildtypes failed to turn pink, and those that did turn pink tended to survive until the end of the observation period (Figure 2B). Partial to full air penetration into the lungs was observed in the deceased mutants that were not stillborn (Supplementary Figure 2A). Together, these results indicate the death of majority of *6qE3^del/+^* pups occurred in the immediate postnatal period due to an inability to oxygenate their blood.

**Figure 2.**
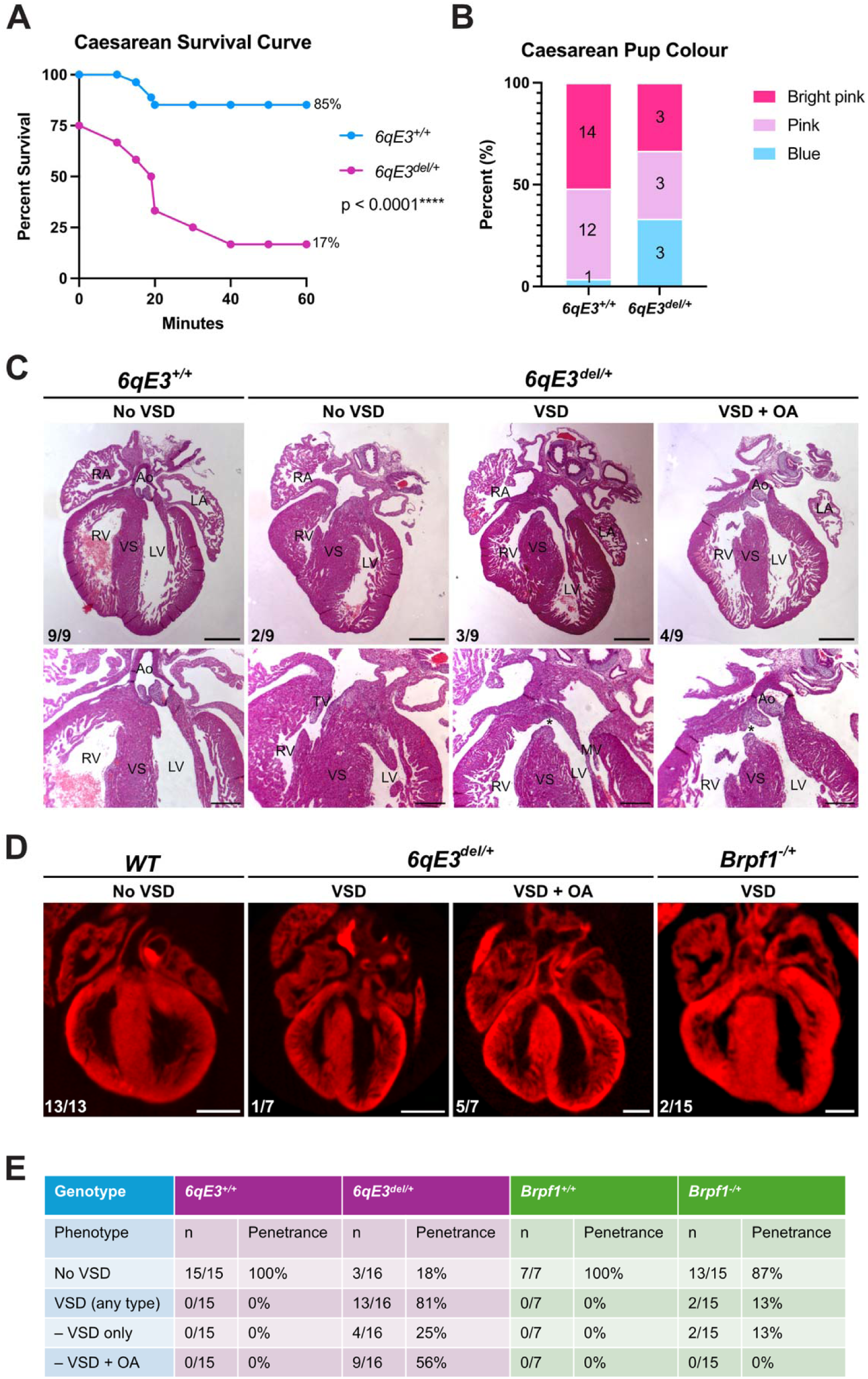
**(A)** E18.5-E19.5 caesarean survival curve where pups were observed over the first hour of life. *6qE3^del/+^* survival is 75% from 0 minutes as stillborn mice were included in the analysis (n = 3 stillborn, all *6qE3^del/+^*). Logrank (Mantel-Cox) test (χ*^2^* = 20.78, df = 1, *6qE3^+/+^*n = 27, *6qE3^del/+^* n = 12, p < 0.0001). **(B)** Distribution of observed pup colour. *6qE3^+/+^* n = 27, *6qE3^del/+^*n = 9 (stillborn excluded). **(C)** Representative H&E sections of E18.5-E19.5 hearts. *6qE3* n = 9 fetuses per genotype. Scale bar for top panel is 0.5 mm, bottom panel is 0.25 mm. Asterisks in bottom panel indicate VSD. Incidence of each phenotype indicated in bottom left corner for each genotype. See supplementary figure 2B for additional replicates. **(D)** Representative image of micro-CT reconstruction of E18.5-E19.5 hearts. Scale bar = 0.5 mm. Incidence of each phenotype indicated in bottom left corner for each genotype. **(E)** Penetrance of VSDs in *6qE3 and Brpf1* fetuses, combining data from both micro-CT and H&E staining. Fisher’s exact t-test of VSD vs. no VSD (*6qE3* p < 0.0001, *Brpf1* p > 0.999). Note that the low incidence of VSDs in *Brpf1^-/+^* relative to *Brpf1^+/+^* did not reach statistical significance, likely reflecting limited statistical power rather than an absence of effect. Abbreviations: VSD = ventricular septal defect, OA = overriding aorta, RA = right atrium, LA = left atrium, Ao = aorta, RV = right ventricle, LV = left ventricle, VS = ventricular septum.

Failure to oxygenate blood after birth can have many underlying causes, including structural defects in the pulmonary or cardiovascular systems [50]. Since congenital heart defects are often seen in 3p25.3 microdeletion patients, we performed hemotoxylin and eosin (H&E) serial sectioning and micro-computed tomography (microCT) on E18.5-E19.5 hearts to identify any structural abnormalities. We found that 81% of *6qE3^del/+^* fetuses had severe perimembranous ventricular septal defects (VSDs), with 69% of these also having an overriding aorta (Figure 2C-E). To understand whether *Brpf1* haploinsufficiency may contribute to this phenotype, we also examined the hearts of E18.5 *Brpf1^-/+^* fetuses and found small perimembranous VSDs in 13% of *Brpf1^-/+^* E18.5 fetuses, indicating that *Brpf1* haploinsufficiency plays a role, but is not the only driver of the VSD phenotype in *6qE3^del/+^* animals, and does not on its own contribute to the additional overriding aorta. Given that lethality was not observed in *Brpf1^/+^* pups, it is likely that VSDs observed in these pups closed during postnatal development or were small enough to not impact survival outcomes.

A range of congenital heart defects including VSDs have been reported in patients with 3p25.3 microdeletion syndrome (25%, 14/57 cases [3, 5, 9, 11, 12, 51–54]) and to a lesser extent, IDDDFP syndrome (17%, 9/53 cases [24, 29, 32]). Our two models recapitulate this VSD phenotype, with much greater severity in the *6qE3^del/+^* model.

### Introduction of the BALB/c genetic background partially rescues perinatal lethality in *C57BL/6-6qE3^del/+^* mice

The neonatal lethality of the *6qE3^del/+^* animals on the C57BL/6 background precluded any experiments on postnatal animals or continuation of the mutant line after the death of the original *C57BL/6*-*6qE3^del/+^*founders. Inbred mouse strains are recessive across all loci and more susceptible to mutations that may impact survival in comparison to a mixed background [55]. The C57BL/6 strain is particularly sensitive to heart defects, with the same mutation in genes such as *CHF1* (cardiovascular basic helix-loop factor 1) leading to 100% penetrant, large VSDs on a C57BL/6 background, whereas 25% of mice have a small VSD on a BALB/c background [56]. Thus, we generated *F1(CB6)*-*6qE3* offspring by outcrossing our *C57BL/6*-*6qE3^del/+^*(*B6-6qE3^del/+^*) founder male with a wildtype BALB/c *(C-6qE3^+/+^)* female and further backcrossed with wildtype BALB/c to produce *N2(C)-6qE3* and *N3(C)-6qE3* offspring (Figure 3A). As survival and incidence of phenotypes reported did not differ between the F1, N2 and N3 generations (Supplementary Figure 3), data are pooled across all generations, annotated collectively as *CB6-6qE3*.

**Figure 3.**
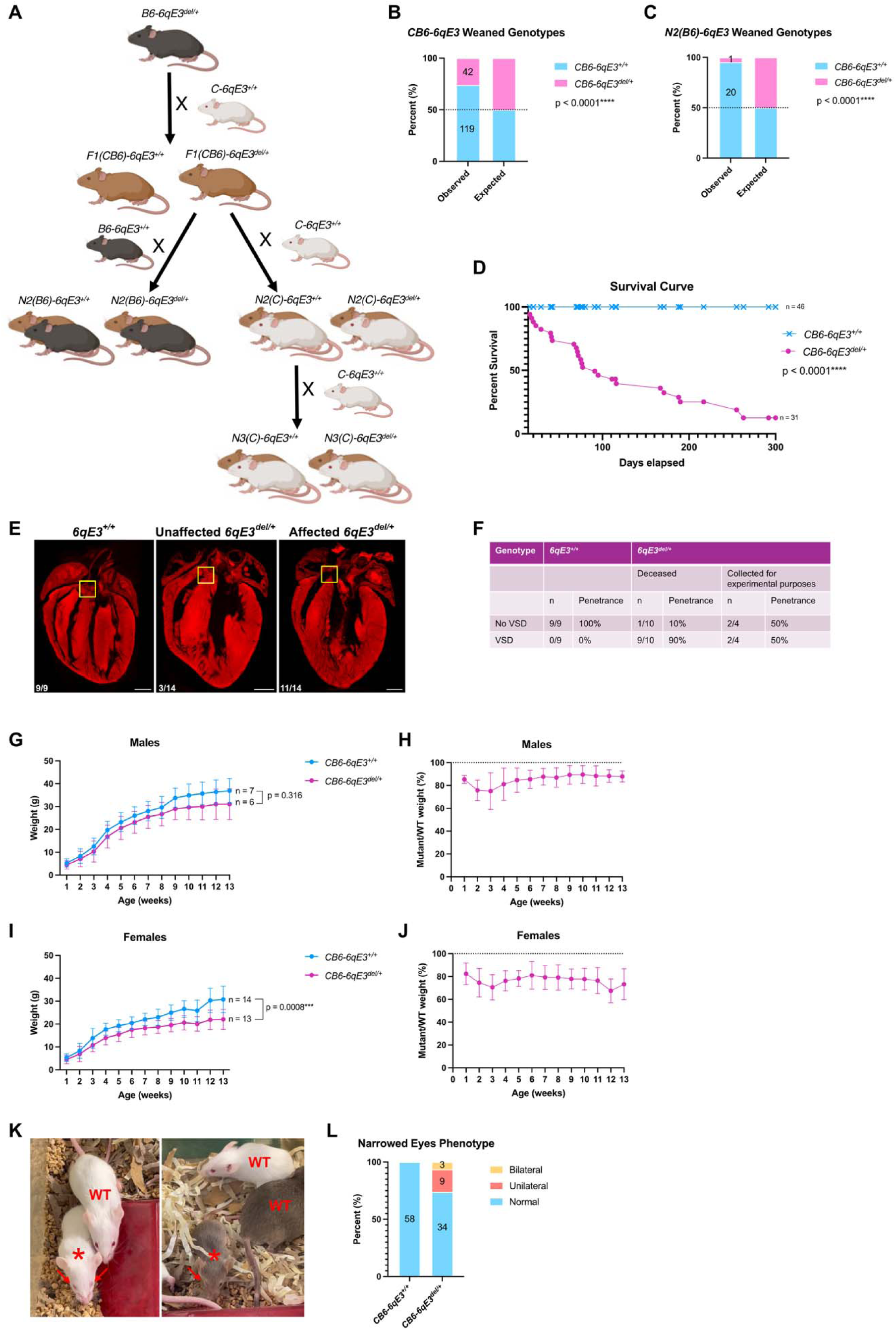
**(A)** Schematic outlining genetic crosses following outcrossing *C57BL/6-6qE3^del/+^* founder to *BALB/c WT*. F1 offspring were then backcrossed to either *C57BL/6 (B6)* or *BALB/c (C)* to make N2 generation mice. Phenotypes did not differ between *F1(CB6)-6qE3, N2(C)-6qE3* and *N3(C)-6qE3*, hence these generations are referred to together collectively as *CB6-6qE3*. **(B)** Genotype distribution of *CB6-6qE3* weaned pups, with observed vs. expected numbers (χ*^2^* = 44.67, df = 1, N = 161, p < 0.0001). **(C)** Genotype distribution of weaned *B6-N2-6qE3* offspring from *CB6-F1-6qE3^del/+^*x *B6-6qE3^+/+^* backcross, with observed vs. expected numbers (χ*^2^* = 17.19, df = 1, N = 21, p < 0.0001). **(D)** Survival curve of *CB6-6qE3* animals. Logrank (Mantel-Cox) test (χ*^2^* = 44.67, df = 1, N = 77, p < 0.0001). X indicates when *CB6-6qE3^+/+^*were collected for experimental purposes. **(E)** Representative images of hearts. VSD indicated by yellow box in affected *6qE3^del/+^*. Penetrance is indicated for whole population analysed in bottom left corner of each image. Scale bar = 1 mm. **(F)** Penetrance of VSD in each genotype, splitting *CB6-6qE3^del/+^* that perished from ones that were collected at 7-8 weeks for experimental purposes. Fisher’s exact t-test (for deceased only) p > 0.0001, Fisher’s exact t-test for total population p = 0.0015. **(G-J)** Growth curves for *CB6-6qE3* **(G)** males and **(I)** females from 1-13 weeks of age, separated by genotype. Two-way repeated measures ANOVA for genotype in males (*CB6-6qE3^+/+^* n = 7, *CB6-6qE3^del/+^*n = 6, p = 0.316) and females (*CB6-6qE3^+/+^* n = 14, *CB6-6qE3^del/+^*n = 13, p = 0.0008). Error bars represent ± 1 SD. **(H)** and **(J)** are the mean values for the weight of each *CB6-6qE3^del/+^* divided by their *CB6-6qE3^+/+^* littermate, expressed as a percentage. Dashed line indicates where weight would be equivalent, below the line means *CB6-6qE3^del/+^*weighed less than their *CB6-6qE3^+/+^* littermate. Error bars represent ± 1 SD. **(K)** Representative images of mice with narrowed eye phenotype, with *CB6-6qE3^+/+^* indicated with ‘WT’ and *CB6-6qE3^del/+^*marked by asterisk. Red arrows point to the affected eyes. **(L)** Quantification of the incidence of the narrowed eye phenotype in *CB6-6qE3*. Fisher’s exact t-test (*CB6-6qE3^+/+^* n = 58, *CB6-6qE3^del/+^*n = 46, p < 0.0001).

In contrast to *B6*-*6qE3^del/+^* animals (Figure 1D), *CB6-6qE3^del/+^* mice were present at weaning, albeit in reduced frequency compared to expected (Figure 3B). Assuming the WT numbers reflect the expected count for both genotypes, 65% of *CB6-6qE3^del/+^*animals are lost prior to weaning, as seen for *B6*-*6qE3^del/+^* animals (Figure 3B). Of the surviving animals, the majority of weaned mutants perished during early adulthood (Figure 3D) after exhibiting signs consistent with heart failure, such as increased respiration, reduced locomotion and altered physical appearance including a hunched posture, ruffled fur and weight loss [57, 58].

When *F1(CB6)*-*6qE3^del/+^* mice were backcrossed to C57BL/6 animals, *N2(B6)-6qE3^del/+^* were almost completely absent at weaning (Figure 3C). Together, this suggests that there is a strong recessive C57BL/6 genetic modifier, or perhaps multiple, that contribute to perinatal lethality of *6qE3^del/+^* on a C57BL/6 background.

Micro-CT was performed on hearts from *CB6-6qE3^del/+^*adult animals that displayed signs consistent with heart failure and *CB6-6qE3^+/+^*littermate controls, revealing that approximately 90% of *CB6-6qE3^del/+^*hearts had a small ventricular septal defect (Figure 3E-F). A small number of otherwise healthy *CB6-6qE3^del/+^* animals were collected at 7-8 weeks of age for experimental purposes and it was found that only 50% of the hearts from this cohort had a small VSD. These observations show that a VSD is present in the majority of adult *6qE3^del/+^* animals even when outbred to BALB/c, however the severity is lessened.

*CB6-6qE3^del/+^* female mice weighed less throughout the first 13 weeks of life compared to *CB6-6qE3^+/+^* animals (Figure 3I), and a similar trend was seen in males (Figure 3G), but this did not reach statistical significance, likely because of the smaller sample size. However, when litter-matching each animal, *CB6-6qE3^del/+^*animals of each sex consistently weighed less than their *CB6-6qE3^+/+^*littermates (Figure 3H,J). A narrowed eye was also observed in approximately 30% of mutants (Figure 3K-L), which may relate to the ptosis seen in 3p25.3 microdeletion syndrome patients [31].

The perinatal lethality seen in *C57BL/6*-*6qE3^del/+^*animals was successfully rescued in approximately 35% of heterozygotes when crossing to BALB/c, though the majority of these mutants still had small VSDs and perish in early adulthood. To further investigate the validity of *6qE3^del/+^*animals as an appropriate model of 3p25.3 microdeletion syndrome, for this study we chose to focus on the *C57BL/6*-*6qE3^del/+^* animals as the genetic system with the greatest phenotypic effect. As *C57BL/6-6qE3^del/+^*resulted in perinatal lethality, we focused on the late gestation period so we could directly compare the two NDD models.

### *6qE3^del/+^* fetuses have sternum defects

3p25.3 microdeletion syndrome patients have a low incidence of spinal malformations such as scoliosis, and pectus excavatum (sunken breastbone) [59, 60]. To investigate whether our models recapitulated these phenotypes, we performed whole skeleton staining of E17.5-E18.5 fetuses with Alcian blue and Alizarin red staining, allowing us to visualise cartilage and ossified bone, respectively.

The skeletons of *6qE3^del/+^* fetuses were broadly phenotypically normal, with the expected number of vertebrae and ribs (Figure 4A-Dii-iii). However, further investigation revealed *6qE3^del/+^*fetuses have subtle sternum defects, such as asymmetric ossification centres (Figure 4A-Div). 80% of *6qE3^del/+^* fetuses had a small sternal foramen, indicative of a development defect as the two sternal bands meet at the midline at approximately E14.5 and begin ossification at E16.5 [61]. *Brpf1* fetuses appeared slightly further into development than *6qE3* fetuses overall based on the increased size and broadly higher levels of ossification, including the sternum, but this did not differ between *Brpf1^+/+^* and *Brpf1^-/+^* fetuses (Figure 4A-Diii). Sternal foramens and asymmetric ossification centres were observed in both *Brpf1^+/+^*and *Brpf1^-/+^* fetuses, though more frequently in *Brpf1^-/+^*fetuses, but this was not statistically significant (Figure 4E). The subtle sternal defects consistently observed in *6qE3^del/+^* and, to a lesser extent, in *Brpf1^-/+^* fetuses, align with previous reports of chest wall deformities including pectus excavatum in 3p25.3 microdeletion syndrome patients, further supporting that this model is reflective of the human disorder.

**Figure 4.**
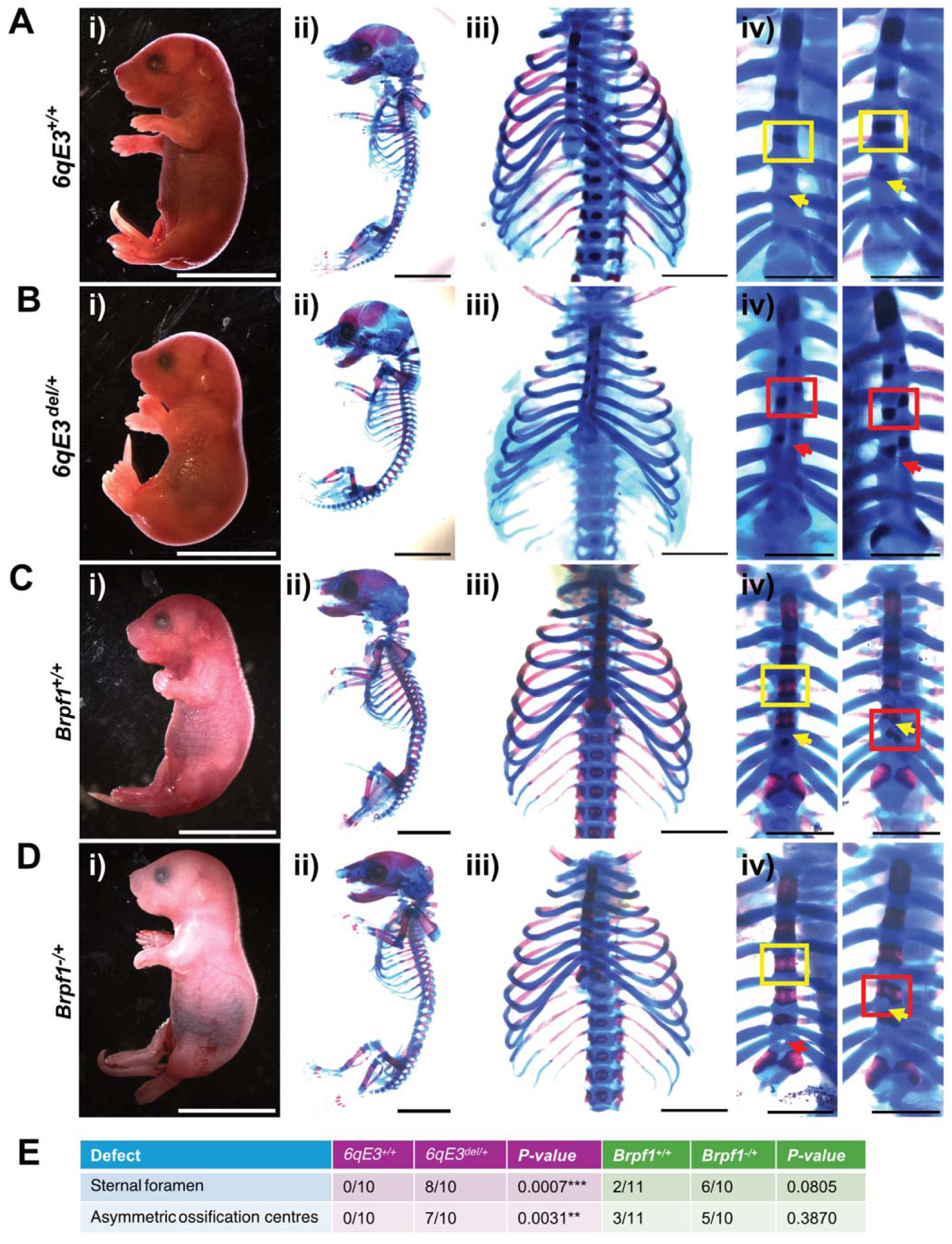
Representative images of **(A)** *6qE3^+/+^,* **(B)** *6qE3^del/+^*, **(C)** *Brpf1^+/+^* and **(D)** *Brpf1^-/+^* E18.5 fetuses. **i.** Gross morphology of fetuses. Scale bar = 1 cm. **ii.** Images of whole skeletons stained with Alcian blue and Alizarin red. Note that extremities such as paws and tails may not be fully stained. Scale bar = 1 cm. **iii.** Representative images of ribs. Scale bar = 2 mm. **iv.** Zoomed in image of the sternum for two fetuses per genotype, with defects indicated with red arrows (sternal foramen) or boxes (asymmetric ossification centres), and normal structures indicated with yellow arrows or boxes. Scale bar = 1 mm. **(E)** Summary table of sternum defects observed across all 4 genotypes, with penetrance and p-values from Fisher’s exact t-test.

### *6qE3^del/+^* and Brpf1^-/+^ fetuses have cranial base defects and other craniofacial abnormalities

To consider whether our new mouse models phenocopied the craniofacial differences observed in patients, we first examined gross morphological features. *6qE3^del/+^* E18.5 fetuses had a subtle decrease in snout length and rounded heads (Figure 5A-B). Conversely, *Brpf1^-/+^* fetuses and adult animals appeared grossly normal.

**Figure 5.**
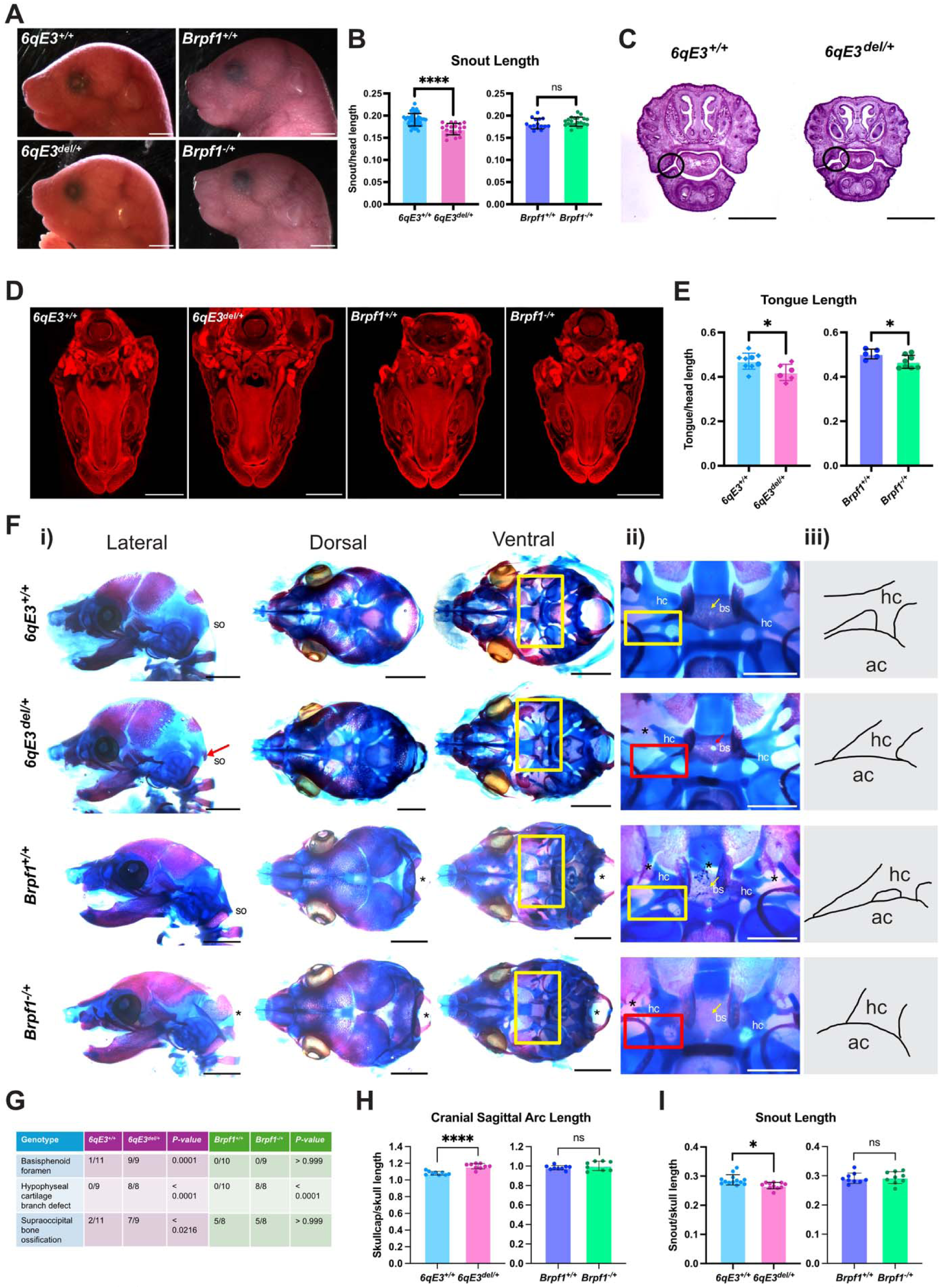
**(A)** Gross facial appearance of E18.5 fetuses. Scale bar = 2 mm. **(B)** Quantification of snout length from gross morphology, normalised to head length. Unpaired t-test two-tailed, *6qE3* (*6qE3^+/+^*n = 43, *6qE3^del/+^* n = 19], p < 0.0001) and *Brpf1* (*Brpf1^+/+^*n = 15, *Brpf1^-/+^* n = 21, p = 0.2397). Error bars represent ± 1 SD. **(C)** Representative H&E stained head sections from *6qE3* E18.5 fetuses. Black circles indicate abnormal shape of maxillary and mandibular processes at the oral commissure. Scale bar = 2 mm. Note that the size of the *6qE3^del/+^* head is markedly smaller than the *6qE3^+/+^*head at this section, which was consistently seen across replicates. **(D)** Representative images of E18.5 tongues at equivalent sections from micro-CT reconstructed scans. Scale bar = 2 mm. **(E)** Quantification of tongue length relative to head length. Diamonds indicate measurements taken from dissections, circles indicate when measured using micro-CT. Unpaired t-test two-tailed, *6qE3* (*6qE3^+/+^* n = 9, *6qE3^del/+^*n = 6, p = 0.0209) and *Brpf1* (*Brpf1^+/+^* n = 5, *Brpf1^-/+^*n = 8, p = 0.0404). Error bars represent ± 1 SD. **(F) i.** Representative images of skulls from the lateral, dorsal and ventral perspective with the jaw removed. Scale bar = 2 mm. Red arrow indicates the premature ossification of supraoccipital bone (so). Asterisks indicate damage to samples that is not a biological effect, whereby skulls for *Brpf1* fetuses caved in during the skeletal staining process. **ii.** Zoomed in ventral view to illustrate sphenoid defects. Hc = hypophyseal cartilage, bs = basisphenoid. Yellow squares or arrows indicate normal structures, red indicates defects. **iii.** Representative traces of the hypophyseal cartilage defect. Ac = auditory capsule. **(G)** Summary of craniofacial defects, penetrance, and the p-value from Fisher’s exact t-test. **(H)** Quantification of cranial sagittal arc length normalised to skull sagittal length. Multiple unpaired t-tests (for all skull measures) using Holm-Šidák correction, *6qE3* (*6qE3^+/+^* n = 9, *6qE3^del/+^*n = 9, p < 0.0001) and *Brpf1* (*Brpf1^+/+^* n = 9, *Brpf1^-/+^*n = 8, p = 0.569). Error bars represent ± 1 SD. **(I)** Quantification of snout length normalised to skull sagittal length. Multiple unpaired t-tests (for all skull measures) using Holm-Šidák correction, *6qE3* (*6qE3^+/+^*n = 13, *6qE3^del/+^* n = 11, p = 0.0100) and *Brpf1* (*Brpf1^+/+^*n = 9, *Brpf1^-/+^* n = 9, p = 0.841). Error bars represent ± 1 SD.

To interrogate internal structures, we performed serial head sectioning with H&E staining of *6qE3^del/+^* fetuses, finding that nasal structures and passageways were unobstructed and grossly equivalent to wildtype controls (Figure 5C). However, subtle structural abnormalities were detected in *6qE3^del/+^* fetuses, such as atypical morphology of the lateral jaw processes at the oral commissure despite a fully closed, normal palate (Figure 5C). Additionally, the tongue appeared to be shorter in *6qE3^del/+^* fetuses, which we confirmed with micro-CT (Figure 5D-E). *Brpf1^-/+^* fetuses did not have the abnormally-shaped lateral jaw processes, but they did have shorter tongues (Figure 5D-E). This may explain why *Brpf1^-/+^*animals weigh less than *Brpf1^+/+^* littermates over the course of postnatal development, as a shortened tongue may impact their ability to feed.

We carried out skeletal staining and found that while sagittal skull length does not change between genotypes, *6qE3^del/+^* fetuses have an increased cranial sagittal arc, indicative of macrocephaly (Figure 5H). Additionally, the nasal bone was similarly decreased in size to what was seen previously from gross morphological evaluation (Figure 5I). *Brpf1^-/+^*fetuses did not share these phenotypes, but there was a very subtle increase in jaw length (Supplementary Figure 4G).

A notable finding from examination of the skulls was the abnormal development of the sphenoid bone, a central anchor for the skull that articulates with almost every other skull bone [62]. *6qE3^del/+^*fetuses had a persistent basisphenoid foramen (Figure 5F-G), consistent with failure of closure of the buccohyposeal canal, a transient structure that typically closes by E12.5 [63]. The hypophyseal cartilage is a transient structure during cranial base development that eventually forms the bulk of the basisphenoid [62]. At E18.5, the hypophyseal cartilage is reduced to branches connecting the basisphenoid and the otic capsule. Both *6qE3^del/+^* and *Brpf1^-/+^* fetuses had abnormal morphology of these branches, where the usual two branches was replaced by a singular thickened branch (Figure 5Fii-iii).

These defects may compromise the structural integrity of the sphenoid, and consequently the skull itself. *6qE3^del/+^* fetuses also had a promiscuous supraoccipital ossification centre, which may further compromise skull base development with increased rigidity. Abnormal sphenoid bone development has been shown to lead to atypical morphology of other skull bones [63, 64], thus potentially explaining the other craniofacial defects observed. Just as patients with 3p25.3 microdeletion syndrome have distinctive facial features, *6qE3^del/+^* fetuses exhibited craniofacial abnormalities, with some overlap seen in *Brpf1^-/+^* fetuses.

### *6qE3^del/+^* fetuses exhibit larger brains, enlarged ventricular spaces and a thinner primary motor cortex

Since our primary interest in these mice is to understand neurodevelopmental disorders, we investigated whether any phenotypes were apparent in the brains of the animals. To examine whether mutants had overt structural brain defects, we performed micro-CT on E18.5-E19.5 whole heads from both colonies. Consistent with the larger head size observed by gross observation (Figure 5A,F), *6qE3^del/+^* fetuses displayed an increase in brain volume between litter-matched pairs from micro-CT imaging of Lugol’s stained brains (Figure 6A-B). Additionally, the *6qE3^del/+^*brains had a larger ventricular volume that also persisted when normalised to total brain volume (Figure 6C-D). In contrast, *Brpf1^-/+^*fetuses did not exhibit these phenotypes (Figure 6C-D).

**Figure 6.**
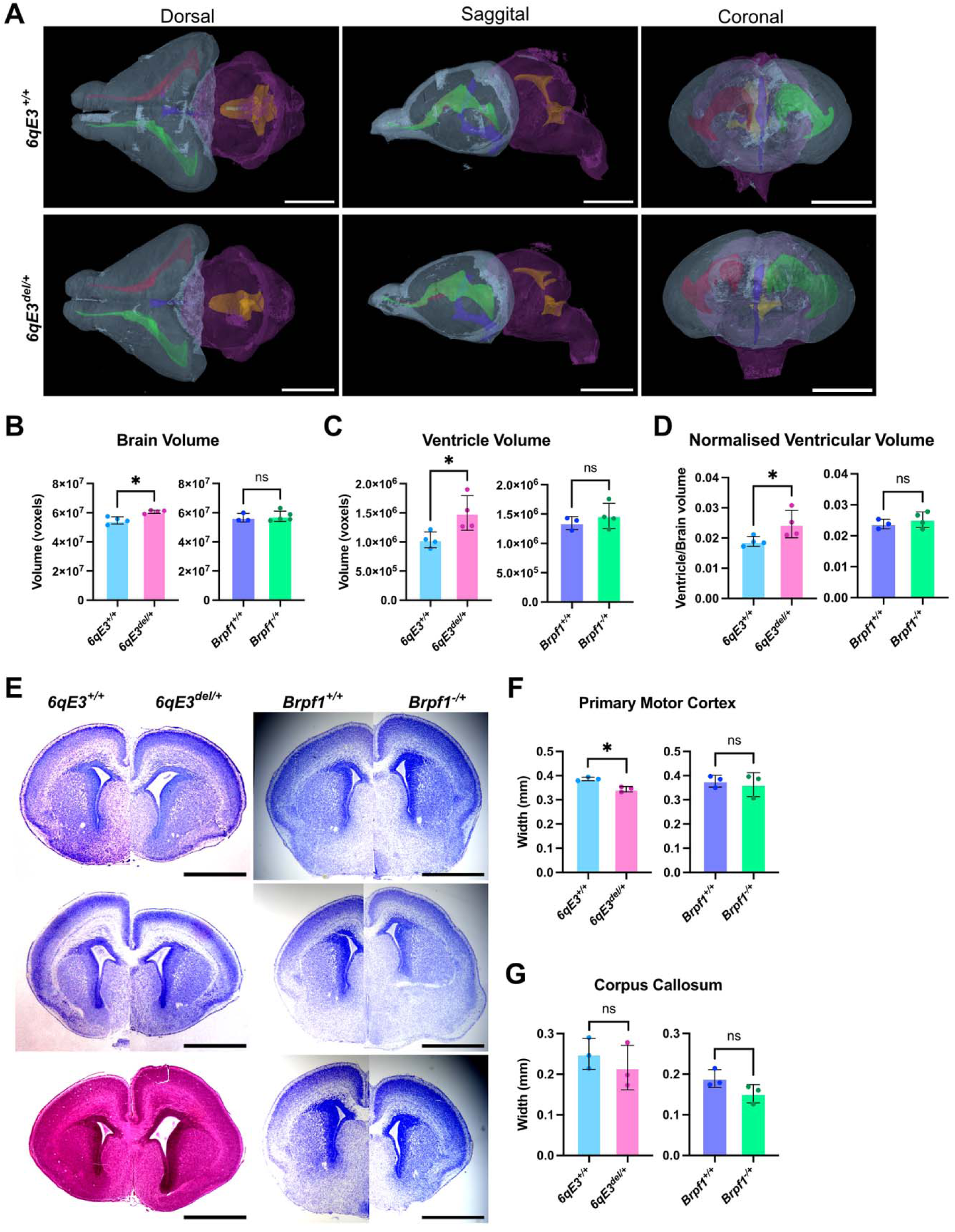
**(A)** Micro-CT 3D-reconstructed images of brains from dorsal, sagittal and coronal planes. Different parts of brain are segmented into different colours: forebrain and olfactory bulbs (blue), hindbrain (purple), lateral ventricles (red and green), 3^rd^ ventricle (dark blue) and 4^th^ ventricle (orange). Note *Brpf1* brains are not shown as they looked equivalent to *6qE3^+/+^*. Scale bar = 1 mm. **(B-D)** Quantification of (B) brain volume (*6qE3* p = 0.011, *Brpf1* p = 0.7043), **(C)** ventricular volume (*6qE3* p = 0.027, *Brpf1* p = 0.6714) and **(D)** ventricular volume normalised to brain volume (*6qE3* p = 0.034, *Brpf1* p = 0.6714). Multiple paired t-tests were performed for *6qE3* (n = 4 per genotype), multiple unpaired t-tests performed for *Brpf1* as the ratio of genotypes was not appropriate for pairing (*Brpf1^+/+^*n = 3, *Brpf1^-/+^* n = 4-5), both with Holm-Šidák correction. Error bars represent ± 1 SD. **(E)** Nissl-stained E18.5 brains. Scale bar = 1 mm. **(F-I)** Quantification of **(F)** brain width (*6qE3* p = 0.277, *Brpf1* p = 0.966), **(G)** brain height (*6qE3* p = 0.708, *Brpf1* p = 0.966), **(H)** primary motor cortex width (*6qE3* p = 0.020, *Brpf1* p = 0.966) and **(I)** corpus callosum width (*6qE3* p = 0.677, *Brpf1* p = 0.369). Multiple unpaired t-tests were performed for both *6qE3* and *Brpf1* (n = 3 per genotype), with Holm-Šidák correction. Error bars represent ± 1 SD.

We further investigated brain structure with Nissl staining of E18.5 brains. Whilst the brain cryoarchitecture generally appeared normal, we identified that the thickness of the anterior primary motor cortex (approximately equivalent to Bregma 2.19 mm) is subtly reduced in *6qE3^del/+^*, but not in *Brpf1^-/+^*, brains compared to littermate controls (Figure 6H). This was not apparent in more medial sections (Supplementary Figure 5D-G). Additionally, cortical layering was unchanged across all genotypes (Supplementary Figure 5H), suggesting that global cytoarchitecture is largely preserved in both *6qE3^del/+^* and *Brpf1^-/+^* fetuses.

Collectively, these data suggest that the structural and cytoarchitectural brain defects observed in *6qE3^del/+^* fetuses are not primarily driven by haploinsufficiency of *Brpf1,* since they are absent in *Brpf1^-/+^* fetuses. To investigate possible changes beyond structural or cytoarchitectural, we asked if there were perturbations at the molecular level.

### H3K23 acetylation is decreased in the primary motor cortex, dentate gyrus and hypothalamus in *Brpf1^-/+^* adult brains

BRPF1 complexes with histone acetyltransferases KAT6A and KAT6B, which are important for laying down H3K23ac and H3K9ac respectively, though functional overlap has been demonstrated [19, 65–67]. Because of this, we asked whether heterozygous *Brpf1* loss, either within the microdeletion interval of *CB6-6qE3^del/+^* animals or in *Brpf1^-/+^*animals, led to a reduction in H3K9ac or H3K23ac. Since our RNA-Seq data from E14.5 NSCs did not reveal large genome-wide changes (Supplementary Table 1-2), nor did fetal brains reveal profound structural and cytoarchitectural changes across both genotypes (Figure 6), we reasoned that the postnatal brain may yield further changes and took advantage of our outbred *CB6-6qE3^del/+^*animals that survive into young adulthood. We performed immunofluorescence microscopy on coronal brain sections and measured acetylation relative to DAPI intensity in specific brain regions for both mouse models, using litter-and sex-matched pairs for pairwise analyses.

*CB6-6qE3^del/+^* brains did not have a statistically significant decrease in H3K9ac or H3K23ac for any brain region tested, nor overall acetylation levels across the entire brain section (Figure 7i-ii, Supplementary Figure 6). *Brpf1^-/+^* samples had no significant decrease in H3K9ac or H3K23ac across the entire brain section (Figure 7Aiii-iv,Eiii-iv). However, H3K23ac was reduced by approximately 26% in the *Brpf1^-/+^* cerebral cortex (Figure 7Biii-iv), specifically the retrosplenial and primary motor cortex (Supplementary Figure 6F). H3K23ac was also reduced by approximately 36% in the *Brpf1^-/+^*hypothalamus (Figure 7Diii-iv). Using NeuN staining to distinguish neurons from other cell types, we confirmed that the H3K23ac decrease is observed in neurons, presumably amongst other cell types, within these regions (Supplementary Figure 6E).

**Figure 7.**
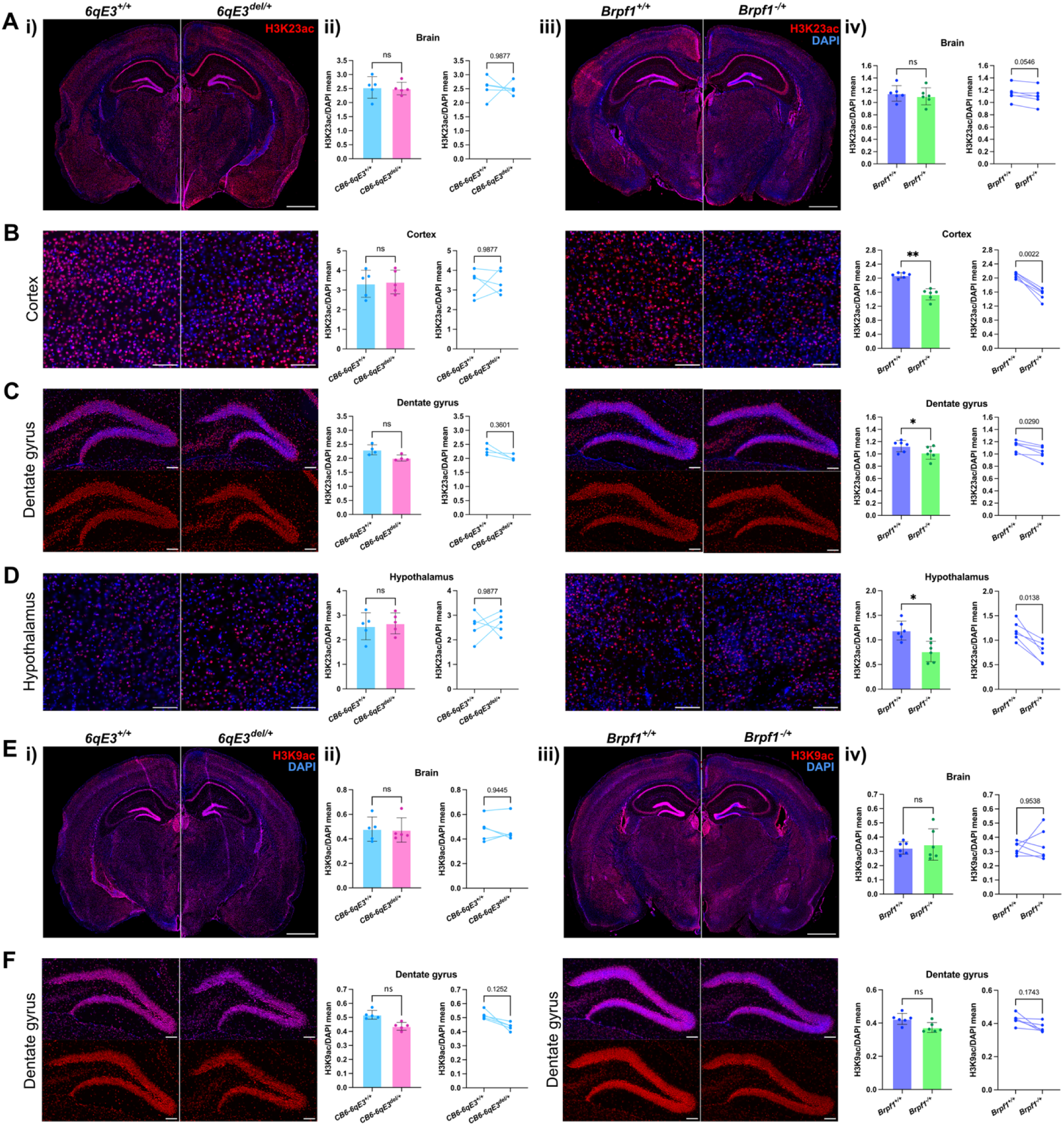
Immunofluorescence of 16 um brain sections from 7-8 week old mice, with **(A-D)** showing H3K23ac in **(A)** whole brains, **(B)** the dorsal cerebral cortex, **(C)** the dentate gyrus, and **(E-F)** showing H3K9ac in **(E)** whole brains and **(F)** the dentate gyrus. Scale bar for whole brains = 1 mm, scale bar for other structures = 100 μm. H3K23ac or H3K9ac appear in red, DAPI appears in blue. **i,iii.** Representative images of i. *CB6-6qE3* (specific generation *N3(C)-6qE3*) and iii. *Brpf1* brains. **ii,iv.** Quantification of acetylation levels in specific areas of the brain, shown as a bar graph (left) and paired line graph (right). Each dot represents an animal (*CB6-6qE3* n = 4-5 per genotype, *Brpf1* N = 6 per genotype), with the mean intensity of the acetylation mark normalised to DAPI. Paired t-tests with Holm-Šidák multiple-testing correction, p-values reported on graphs. Error bars represent ± 1 SD.

Interestingly, both *CB6-6qE3^del/+^* and *Brpf1^-/+^*brains showed a subtle decrease in both H3K9ac and H3K23ac in the dentate gyrus that was consistent across all pairs, though only H3K23ac in the *Brpf1^-/+^*dentate gyrus was statistically significant after multiple testing correction was applied (Holm-Šidák correction, p < 0.05, Figure 7C,F). Given the dentate gyrus is important for memory formation and learning [68], the reduction in acetylation in *Brpf1^-/+^*samples suggests the dentate gyrus may have impaired functioning, consistent with intellectual disability.

Overall, this shows that *Brpf1* haploinsufficiency results in H3K23ac reduction in the primary motor cortex, hypothalamus and dentate gyrus, suggesting reduced KAT6A/B activity.

## Discussion

In this study, through creation of two new mouse models of NDDs, we have demonstrated that *Brpf1* haploinsufficiency is contributing to a select few of the phenotypes seen in *6qE3^del/+^* animals, such as shortened tongues, congenital heart defects and subtle cranial base defects. However, *6qE3^del/+^* phenotypes tended to be more severe, which is consistent with the human disorders. Additionally, *6qE3^del/+^* fetuses had phenotypes not seen in *Brpf1^-/+^* fetuses, such as rounded heads and shortened snouts which are suggestive of the distinctive craniofacial features observed in human patients. This suggests that other genes contained within the deletion interval are contributing to the phenotypes we observed.

The ventricular septal defects seen in *6qE3^del/+^* fetuses were more penetrant and severe than in *Brpf1^-/+^* fetuses, suggesting that other genes contained within this deletion interval may also be contributing to this phenotype. Heterozygous variants in *SETD5* (SET domain containing 5), another gene included in our deletion interval, leads to the NDD SETD5-related syndrome. *Setd5^-/+^* mice present with VSDs and a low incidence of double outlet right ventricle [69]. Thus, *Setd5* loss is likely contributing to the increased severity of VSDs seen in *6qE3^del/+^*compared to *Brpf1^-/+^* animals.

Our work investigated structural and cytoarchitectural changes in *6qE3^del/+^* and *Brpf1^-/+^* fetuses. Previous reports have demonstrated that Brpf1 has an important role in brain electrophysiology in mice, where a 50% reduction in *Brpf1* using *shBrpf1* leads to the decreased frequency of miniature excitatory postsynaptic currents in hippocampal neurons [43]. Additionally, disruption of another gene contained in the *6qE3^del^* region, *LHFPL4* (LHFPL tetraspan subfamily member 4), has been linked to abnormal brain functioning. *Lhfpl4* loss leads to reduction of inhibitory transmission at GABA-A receptors, with *Lhfpl4^-/-^* mice experiencing motor problems and premature death [70, 71]. Thus, further work with functional brain and behavioural phenotyping of *6qE3^del/+^* on the BALB/c background could provide functional insights in addition to the structural defects described here.

*Brpf1* is crucial for dentate gyrus formation, with forebrain-specific inactivation showing mice have an underdeveloped suprapyramidal blade and complete loss of the infrapyramidal blade of the hippocampus [40]. We observed decreased H3K23ac in the dentate gyrus of *Brpf1^-/+^* animals, which aligns with previous reports of decreased H3K23ac in the dorsal cortex of forebrain-specific *Brpf1*-knockout pups [29]. Along with previous behavioural testing that has demonstrated hippocampal-based learning and memory deficits in *Brpf1^-/+^* mice [41], together this perhaps suggests acetylation dysregulation as the contributing molecular mechanism of these phenotypes. We also saw the suggestion of a decrease in H3K23ac in *CB6-6qE3^del/+^* animals, and H3K9ac in both mouse models. Since we assayed H3K9ac and H3K23ac by immunofluorescence, the results are a measure of these marks at the global level. It is plausible that the effect of *Brpf1* haploinsufficiency on acetylation levels in both *Brpf1^-/+^*and *6qE3^del/+^* brains and other tissues are generally more subtle than can be detected at a genome-wide level and will be better detected using genomics-based approaches such as CutNTag to identify locus-specific changes.

BRPF1’s histone acetyltransferase complex members KAT6A and KAT6B have both shown decreased acetylation in heterozygous mouse models. In the adult mouse brain, *Kat6a^-/+^* mice have decreased H3K23ac and *Kat6b^-/+^*mice have decreased H3K9ac, but upon treatment with an acetyl donor, acetyl-L-carnitine (ALCAR), *Kat6a^-/+^* and *Kat6b^-/+^*mice have comparable acetylation levels to their WT counterparts [18, 20]. This has further been shown to lead to a rescue in neurodevelopmental behaviours in both models, such as learning deficits. Thus, we propose that treatment with ALCAR may lead to amelioration of the phenotypes we have described in *6qE3^del/+^* and *Brpf1^-/+^* mice.

Sphenoid defects were seen in both *6qE3^del/+^* and *Brpf1*^-/+^ animals, with both sharing abnormal morphology of the remnant hypophyseal cartilage, and *6qE3^del/+^* having a persistent basisphenoid foramen. As the primary anchor in the skull, minor aberrations of the sphenoid compromise cranial stability, which has been shown to lead to significant craniofacial abnormalities, and often do not occur in isolation [63, 72]. Similar to *6qE3^del/+^*, *Shh^+/-^*mice have a basisphenoid foramen at E18.5, which was linked with a rounded curvature of the skull that worsened as mice aged [63]. Thus, the rounded head phenotype we observed could be a secondary consequence to these subtle sphenoid defects. Cranial instability can also lead to motor and sensory deficits and vision issues [73], both of which align with the human syndromes [11, 29]. The thickened remnant hypophyseal cartilage branch we saw in *6qE3^del/+^* also further suggests that earlier processes in sphenoid development were dysregulated.

While the phenotypes we observed in our 3p25.3 microdeletion syndrome mouse model are not solely driven by *Brpf1*, it is useful to delineate *Brpf1*’s role in this disorder and gain further insights that may inform treatment avenues in future work. By generating a mouse model for 3p25.3 microdeletion syndrome, we were able to faithfully recapitulate many phenotypes seen in human patients, providing an appropriate model that can be used to elucidate the underlying basis of disease, and to potentially develop therapeutic treatments.

## Acknowledgements

The authors thank Lachlan Whitehead for micro-CT acquisition and teaching the analysis pipeline, all members of the WEHI Animal Facility for expert animal care, particularly Ani Lyristis, Louise Spencer, Carmen Gatt, Sasanka Wijethunga and Kevin Tran, the WEHI Advanced Histotechnology Facility for histological processing and slide scanning (Ellen Tsui and Lei Liu), the WEHI Advanced Genomics Facility for sequencing services, Anna Le Fevre for expert clinician insights, and the entire Blewitt laboratory (Boya Zhang, Ruifeng Hu, Kelsey Breslin, Tamara Cameron, Megan Iminitoff, Caleb Chew, Henry Bell and Andres Tapia del Fierro) for invaluable discussions and assistance.

This research was supported by the Commonwealth through an Australian Government Research Training Program Scholarship [DOI: https://doi.org/10.82133/C42F-K220]; the Pamela and Lorenzo Galli Charitable Trust; the Brain M Davis Charitable Trust; the Walter and Eliza Hall Biomedical Research Institute PhD Student Top Up Scholarship; NHMRC Investigator grant (MEB); Al and Val Rosenstrauss Fellowship (HKV) and NHMRC IRIISS and Victorian State Government Operational Infrastructure Support. The generation of *C57BL/6*-*6qE3^del/+^*and *C57BL/6*-*Brpf1^-/+^* mice used in this study was supported by Phenomics Australia and the Australian Government through the National Collaborative Research Infrastructure Strategy (NCRIS) program.

## Authorship Statement

SM co-designed and performed the experiments, carried out data analyses, generated the figures, and drafted the manuscript. MB and HKV conceived the project, supervised the work, designed experiments, guided data interpretation and edited the manuscript. HV and SE contributed to the dissection of *6qE3* E18.5 skeletons and *CB6-6qE3* brains. AK (Kueh) created the mouse lines. AK (Keniry) performed the RNA-sequencing alignment, quality control, and differential expression analysis. All authors approve of the content of the manuscript and agree to be held accountable for the work.

