## Supplementary Data for "A novel mouse model recapitulates features of the neurodevelopmental disorder 3p25.3 microdeletion syndrome"

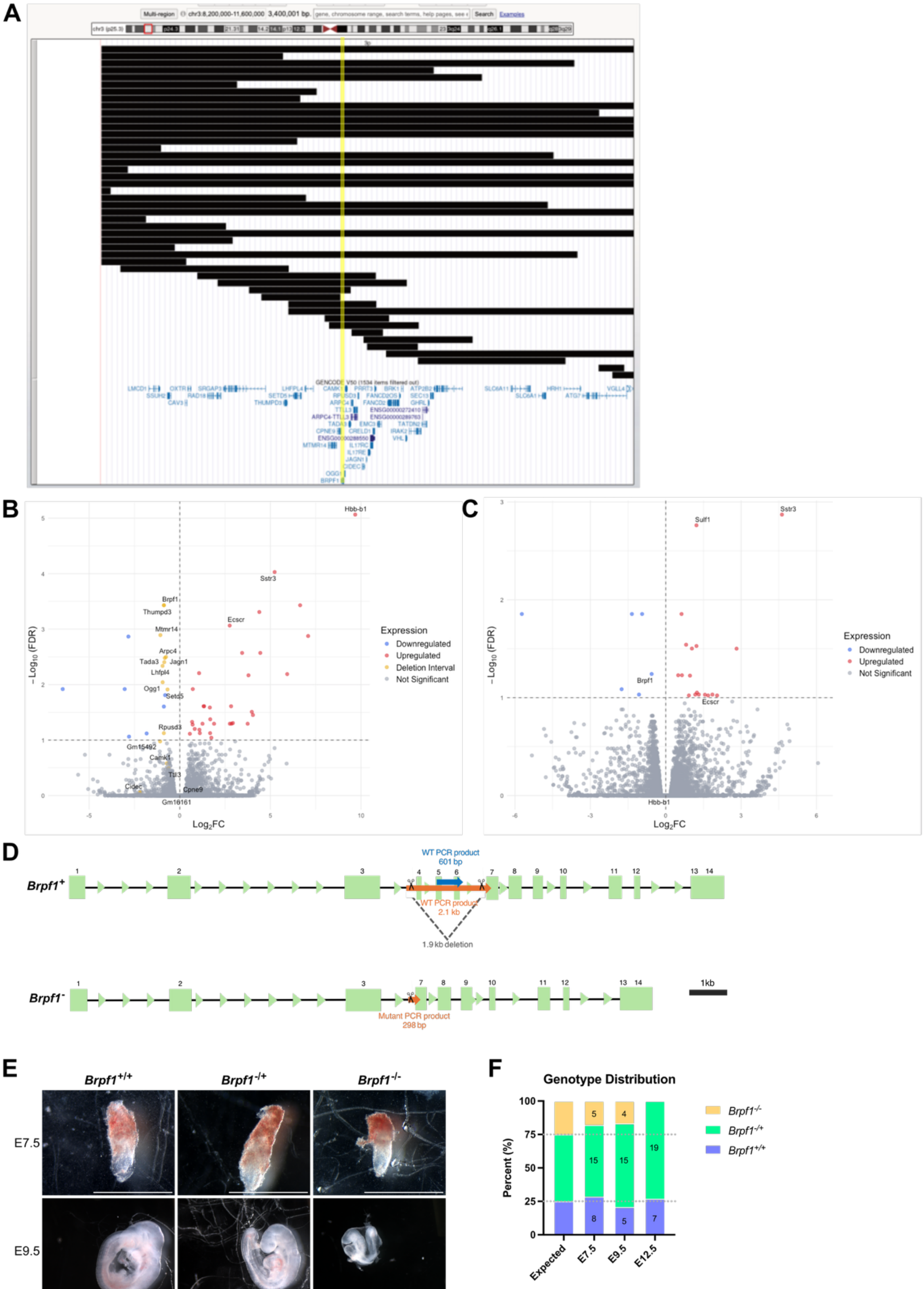


G

| | $Brpf1^{+/+}$ | | $Brpf1^{-/+}$ | | $Brpf1^{-/-}$ | | Total (*n*) |
|---|---|---|---|---|---|---|---|
| | *n* | % | *n* | % | *n* | % | |
| E7.5 | 8 | 28.6 | 15 | 53.6 | 5 | 17.9 | 28 |
| E9.5 | 5 | 20.8 | 15 | 62.5 | 4 | 16.7 | 24 |
| E12.5 | 7 | 26.92 | 19 | 73.08 | 0 | 0 | 26 |

**Supplementary figure 1. (A)** 3p25.3 microdeletion syndrome patients deletion breakpoints (from ClinVar) mapped onto the UCSC genome browser, with the viewing window showing the entire 3p25.3 locus. The yellow rectangle highlights the location of BRPF1, which is heterozygously deleted in 50% (24/48) of patients from the ClinVar database. Note that older published cases do not have specified breakpoints, and therefore aren't included. **(B-C)** Volcano plot of differential gene expression from RNA-Seq of E14.5 NSCs for **(B)** *6qE3* and **(C)** *Brpf1*. Each point represents a gene, plotted by $\log_2$ fold change (FC) against $-\log_{10}$(FDR), with upregulated genes in red, downregulated genes in blue, and yellow genes indicating those included in the $\mathit{6qE3}^{del}$ deletion interval for (B). **(D)** Schematic representation of $\mathit{Brpf1}^{-/+}$ mouse line. Mice are heterozygous for a 1.9-kb deletion in *Brpf1*. Boxes represent exons. Scissors mark the sgRNA cut sites. Orange represents the size of the PCR products when genotyping ($\mathit{Brpf1}^{+}$ band is 2.1 kb, $\mathit{Brpf1}^{-}$ band is 310 bp). Blue represents the WT PCR product that is made by amplifying a region fully inside the deleted segment. **(E-G)** Embryonic litters collected from $\mathit{Brpf1}^{-/+}$ crosses. **(E)** Representative images of each genotype at different embryonic stages. $\mathit{Brpf1}^{-/-}$ was not present at E12.5. Scale bar = 1 mm. **(F)** Graphical representation of genotype distribution. Numbers indicate animals per genotype. Dotted lines indicate expected distribution. **(G)** Table with genotype distribution from litters collected.

**Supplementary Table 1.** Significant hits from RNA sequencing of *6qE3* E14.5 NSCs using FDR correction (FDR < 0.1). $Log_2$ fold change values are generated by subtracting *Brpf1*$^{-/+}$ from WT (negative values indicate upregulation, positive values indicate downregulation).

| Gene | Chr | Start | End | P-value | FDR | Log2 Fold Change |
|---|---|---|---|---|---|---|
| Hbb-b1 | 7 | 103826534 | 103828096 | 3.31E-10 | 8.61E-06 | -9.679896 |
| Hbb-b2 | 7 | 103812524 | 103813996 | 4.61E-07 | 0.00133131 | -7.084 |
| Hba-a1 | 11 | 32283511 | 32284465 | 6.66E-08 | 3.71E-04 | -6.6415424 |
| Gdnf | 15 | 7811011 | 7837575 | 4.48E-06 | 0.00646616 | -5.9277124 |
| Sstr3 | 15 | 78537015 | 78544345 | 7.20E-09 | 9.35E-05 | -5.2361245 |
| Mmp3 | 9 | 7445822 | 7455972 | 1.22E-06 | 0.00268574 | -4.441461 |
| Pdyn | 2 | 129686565 | 129699844 | 1.14E-07 | 4.91E-04 | -4.3890343 |
| Rxfp2 | 5 | 150018675 | 150082184 | 4.35E-05 | 0.03529614 | -4.0361047 |
| Vstm2l | 2 | 157914653 | 157944719 | 3.70E-05 | 0.03098885 | -3.9749517 |
| Fxyd1 | 7 | 31051681 | 31055656 | 4.97E-06 | 0.00679481 | -3.7908037 |
| Mip | 10 | 128225838 | 128231812 | 7.16E-05 | 0.05077014 | -3.7439332 |
| 1700023F06Rik | 11 | 103198946 | 103208558 | 1.24E-06 | 0.00268574 | -3.4410539 |
| Crh | 3 | 19693401 | 19695396 | 6.65E-05 | 0.04934038 | -2.9294736 |
| Gm13152 | 4 | 147492417 | 147513486 | 7.82E-05 | 0.05077014 | -2.8529387 |
| Hephl1 | 9 | 15051841 | 15112108 | 2.65E-05 | 0.02452791 | -2.8316576 |
| Pla2g5 | 4 | 138799244 | 138863482 | 7.54E-05 | 0.05077014 | -2.7638311 |
| Ecscr | 18 | 35713086 | 35722356 | 2.32E-07 | 8.62E-04 | -2.7555873 |
| Wisp1 | 15 | 66891320 | 66923201 | 7.67E-05 | 0.05077014 | -1.8918426 |
| Car10 | 11 | 93098028 | 93601749 | 1.75E-04 | 0.09077101 | -1.740895 |

| | | | | | | |
|---|---|---|---|---|---|---|
| Il1r2 | 1 | 40084768 | 40125219 | 5.36E-05 | 0.04219388 | -1.6817958 |
| Tmem59l | 8 | 70483876 | 70487358 | 1.33E-04 | 0.0750374 | -1.6742696 |
| Ppl | 16 | 5086291 | 5132481 | 2.99E-05 | 0.02589414 | -1.6600481 |
| Thbs2 | 17 | 14665500 | 14694262 | 2.60E-05 | 0.02452791 | -1.3446877 |
| Pdgfd | 9 | 6168612 | 6378843 | 2.64E-05 | 0.02452791 | -1.317904 |
| Clcf1 | 19 | 4214238 | 4223490 | 7.74E-05 | 0.05077014 | -1.2526364 |
| Akr1b8 | 6 | 34354119 | 34368463 | 1.31E-04 | 0.0750374 | -1.1095876 |
| Fcgrt | 7 | 45092993 | 45103822 | 1.08E-04 | 0.06514874 | -1.0840938 |
| Timp1 | X | 20870166 | 20874735 | 4.07E-06 | 0.00622242 | -1.0653349 |
| Tmem108 | 9 | 103482936 | 103761837 | 1.02E-04 | 0.06300643 | -0.8243713 |
| Scn1b | 7 | 31116525 | 31126945 | 1.03E-05 | 0.01203622 | -0.7190116 |
| Ildr2 | 1 | 166254139 | 166316823 | 8.32E-05 | 0.05267195 | -0.7109358 |
| Ndufa3 | 7 | 3617373 | 3620163 | 6.15E-05 | 0.04699423 | -0.6819146 |
| Pygm | 19 | 6384399 | 6398459 | 1.42E-04 | 0.07697146 | -0.5562593 |
| Setd5 | 6 | 113077365 | 113153435 | 1.13E-05 | 0.01222755 | 0.67623353 |
| Arpc4 | 6 | 113378113 | 113390448 | 1.59E-06 | 0.00317898 | 0.76228386 |
| Pcx | 19 | 4510472 | 4621752 | 1.49E-05 | 0.01549003 | 0.80834204 |
| Tada3 | 6 | 113366025 | 113377883 | 1.79E-06 | 0.00331467 | 0.82695043 |
| Jagn1 | 6 | 113442569 | 113448229 | 2.28E-06 | 0.00394603 | 0.8546462 |
| Brpf1 | 6 | 113307181 | 113324857 | 7.14E-08 | 3.71E-04 | 0.87124527 |
| Igf2bp2 | 16 | 22059009 | 22163299 | 2.78E-05 | 0.02486814 | 0.87798315 |
| Thumpd3 | 6 | 113046225 | 113068273 | 4.78E-08 | 3.71E-04 | 0.87824404 |
| Rpusd3 | 6 | 113415319 | 113419348 | 1.28E-04 | 0.0750374 | 0.8789281 |

| | | | | | | |
|---|---|---|---|---|---|---|
| Ogg1 | 6 | 113326972 | 113335068 | 6.98E-06 | 0.00906804 | 0.9417966 |
| Lhfpl4 | 6 | 113168090 | 113195384 | 2.84E-06 | 0.00460371 | 0.94763446 |
| Mtmr14 | 6 | 113237843 | 113281387 | 3.94E-07 | 0.00127997 | 1.0704457 |
| Vmn1r90 | 7 | 14530654 | 14623066 | 1.38E-04 | 0.07597966 | 1.826922 |
| Gm239 | 10 | 116776545 | 116896879 | 1.64E-04 | 0.08671929 | 2.7997093 |
| Slc38a4 | 15 | 96994823 | 97055956 | 5.24E-07 | 0.0013617 | 2.8351634 |
| Art1 | 7 | 102101743 | 102111148 | 1.07E-05 | 0.01203622 | 3.0375679 |
| Xlr4a | X | 73074345 | 73082478 | 1.06E-05 | 0.01203622 | 6.4569607 |

**Supplementary Table 2.** Significant hits from RNA sequencing of *Brpf1* E14.5 NSCs using FDR correction (FDR < 0.1). $\text{Log}_2$ fold change values are generated by subtracting *6qE3*$^{del/+}$ from WT (negative values indicate upregulation, positive values indicate downregulation).

| Gene | Chr | Start | End | P-value | FDR | $\text{Log}_2$ Fold Change |
|---|---|---|---|---|---|---|
| Sstr3 | 15 | 78537015 | 78544345 | 5.18E-08 | 0.00134559 | -4.624357 |
| Sulf1 | 1 | 12692430 | 12860371 | 1.33E-07 | 0.00173114 | -1.220738 |
| Vmn1r32 | 6 | 66552183 | 66559708 | 1.64E-06 | 0.01395722 | 5.72674 |
| Pdgfra | 5 | 75152291 | 75198204 | 2.62E-06 | 0.01395722 | 0.94123447 |
| Pex5l | 3 | 32949408 | 33143247 | 2.84E-06 | 0.01395722 | -0.6293191 |
| Fzd9 | 5 | 135248938 | 135251230 | 3.23E-06 | 0.01395722 | 1.3459411 |
| Il10rb | 16 | 91406164 | 91425834 | 7.75E-06 | 0.02876112 | -0.810141 |
| Cd34 | 1 | 194938821 | 194961292 | 9.14E-06 | 0.02965469 | -1.2221241 |
| Il6ra | 3 | 89869324 | 89913162 | 1.09E-05 | 0.03147805 | -1.0416093 |
| G630090E17Rik | 10 | 39946907 | 39960153 | 1.22E-05 | 0.03156588 | -2.8178897 |
| Brpf1 | 6 | 113307181 | 113324857 | 2.43E-05 | 0.05728645 | 0.56058985 |
| Sirpa | 2 | 129592835 | 129632228 | 2.82E-05 | 0.05897637 | -0.4956498 |
| Adamts14 | 10 | 61197112 | 61273438 | 2.95E-05 | 0.05897637 | -0.6456966 |
| Cdkn2a | 4 | 89274473 | 89294653 | 3.21E-05 | 0.05950875 | -0.9636533 |
| Scrg1 | 8 | 57455923 | 57477585 | 4.73E-05 | 0.0818927 | 1.7555779 |
| C1qtnf1 | 11 | 118428203 | 118449963 | 5.47E-05 | 0.08869284 | -1.2406318 |
| Aldh1a7 | 19 | 20692953 | 20727562 | 6.30E-05 | 0.09295233 | -1.8559436 |
| Slc7a3 | X | 101079210 | 101086020 | 7.00E-05 | 0.09295233 | 1.0660173 |
| Abcg1 | 17 | 31057698 | 31115777 | 7.15E-05 | 0.09295233 | -1.1746058 |

| | | | | | | |
|---|---|---|---|---|---|---|
| Ccdc3 | 2 | 5137776 | 5230878 | 7.16E-05 | 0.09295233 | -1.31646 |
| BC007180 | 3 | 154762157 | 154767787 | 7.55E-05 | 0.0933281 | -1.5602876 |
| Ecscr | 18 | 35713086 | 35722356 | 8.22E-05 | 0.09486653 | -2.0420115 |
| Lama3 | 18 | 12334024 | 12583013 | 8.65E-05 | 0.09486653 | -1.687388 |
| 9930013L23Rik | 7 | 83932857 | 84086502 | 8.77E-05 | 0.09486653 | -0.9197363 |

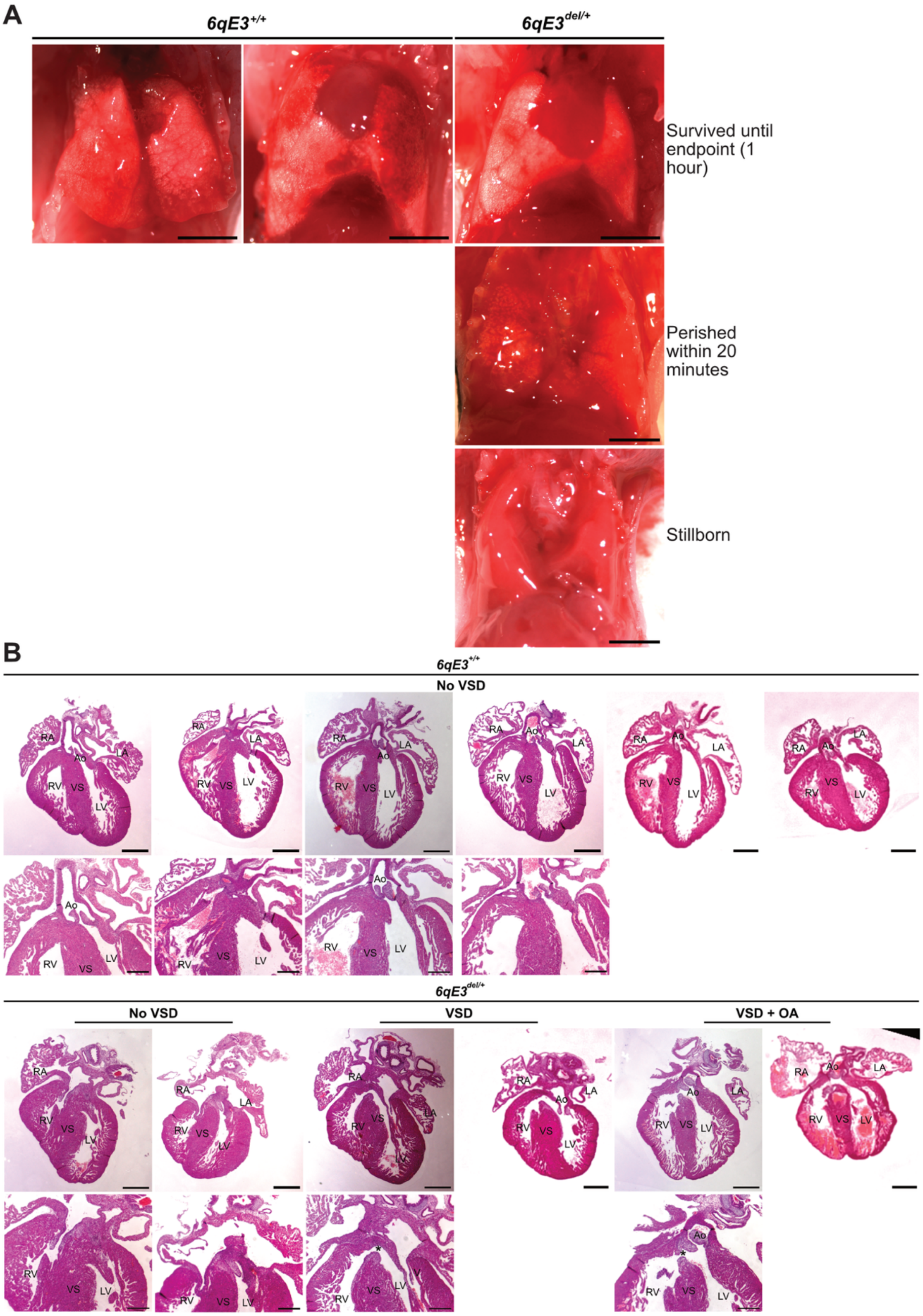


**Supplementary figure 2. (A)** Representative images of lungs from caesarean experiments, showing air penetration in lungs from pups with differential survivability.

Scale bar = 2 mm. **(B)** Additional replicates for H&E serial heart sections of E18.5-E19.5 hearts (data shown in Figure 2C). VSD = ventricular septal defect, OA = overriding aorta, RA = right atrium, LA = left atrium, Ao = aorta, RV = right ventricle, LV = left ventricle, VS = ventricular septum. N = 9 per genotype. Asterisks indicate VSD. Scale bar for whole heart is 0.5 mm, zoomed in region is 0.25 mm.

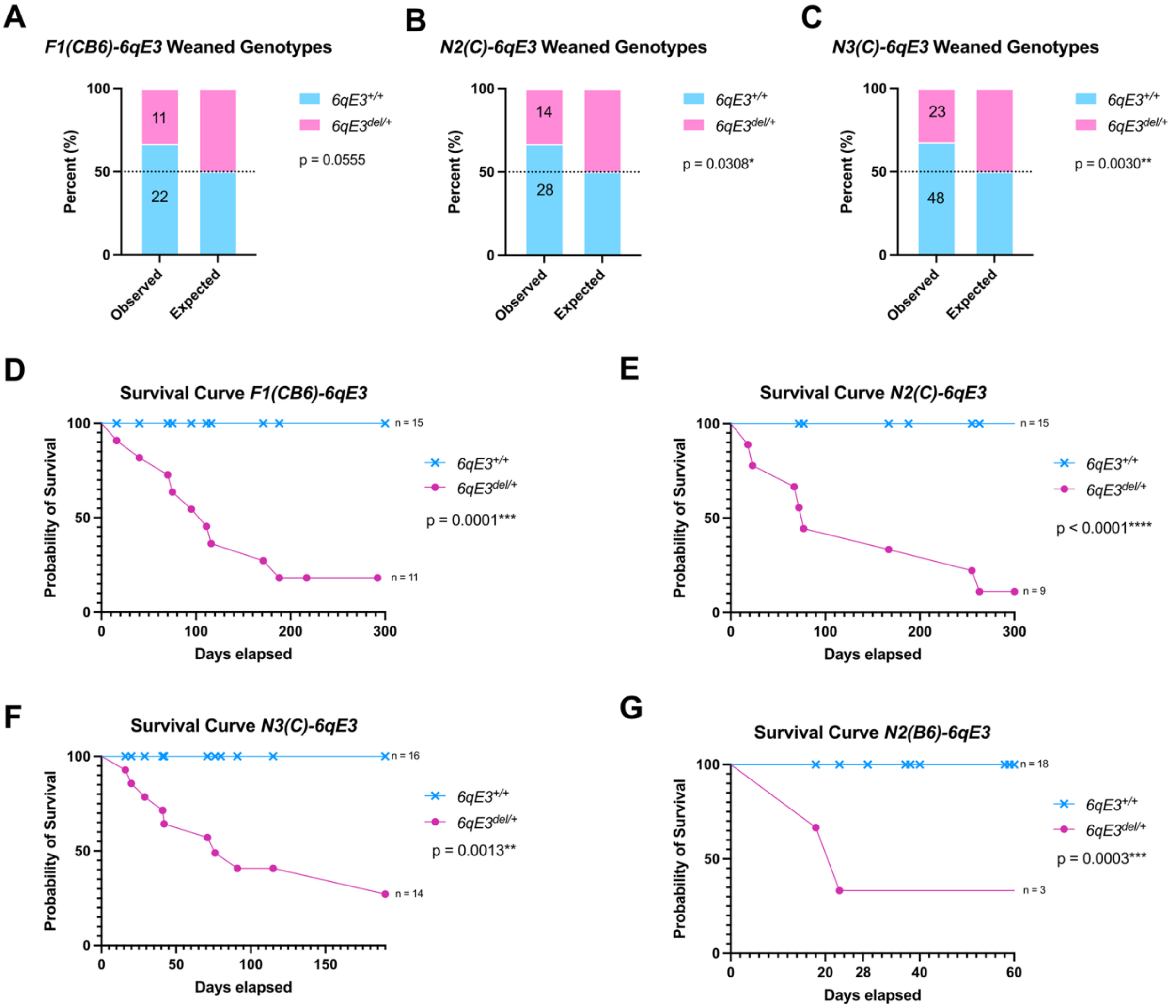


**Supplementary Figure 3. (A-C)** Weaned genotype distributions for **(A)** *F1(CB6)* generation with observed vs. expected numbers ($\chi^2$ = 3.667, df = 1, N = 33, p = 0.0555), **(B)** *N2(C)* generation ($\chi^2$ = 4.667, df = 1, N = 42, p = 0.0308), and **(C)** *N3(C)* generation ($\chi^2$ = 8.803, df = 1, N = 71, p = 0.0030). Note Figure 3B in main text shows **(A-C)** combined as frequencies do not differ across generations. **(D-G)** *CB6-6qE3* survival curves for **(D)** *F1(CB6)* generation, Logrank (Mantel-Cox) test ($\chi^2$ = 14.90, df = 1, p = 0.0001), **(E)** *N2(C)* generation, Logrank (Mantel-Cox) test ($\chi^2$ = 15.25, df = 1, p < 0.0001), **(F)** *N3(C)* generation, Logrank (Mantel-Cox) test ($\chi^2$ = 10.34, df = 1, p = 0.0013) and **(G)** *N2(B6)* generation, Logrank (Mantel-Cox) test ($\chi^2$ = 13.37, df = 1, p = 0.0003).

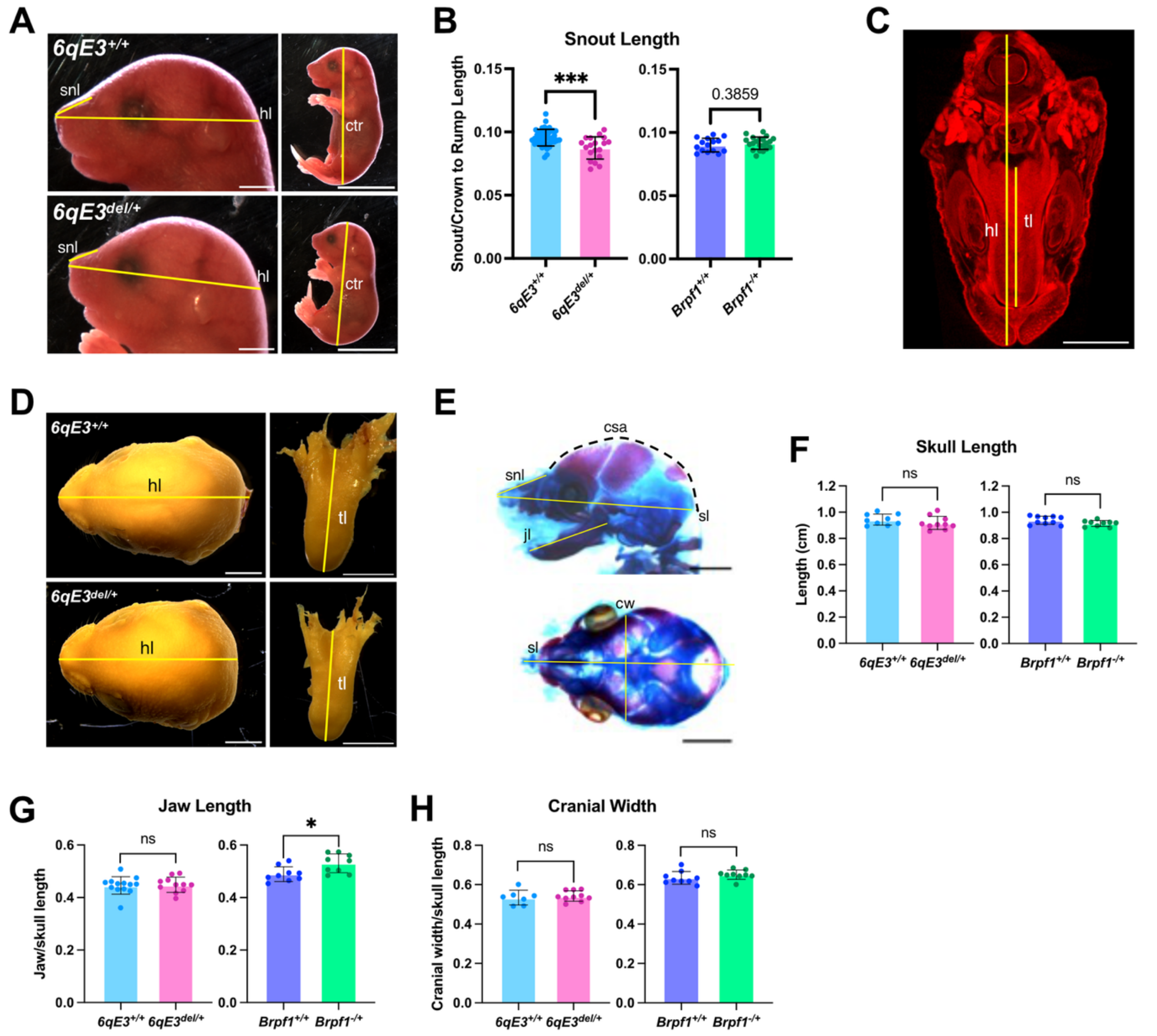


**Supplementary Figure 4. (A)** Measurement references for E18.5 fetuses gross morphology (data shown in Figure 5B). Left scale bar = 2 mm, right is 1 cm. **(B)** Quantification of gross snout length normalised to crown to rump length. Unpaired t-test, *6qE3* (*6qE3*$^{+/+}$ n = 43, *6qE3*$^{del/+}$ n = 19, p < 0.0001) and *Brpf1* (*Brpf1*$^{+/+}$ n = 15, *Brpf1*$^{-/+}$ n = 21, p = 0.2397). Error bars represent ± 1 SD. **(C)** Measurement references for micro-CT tongues (data shown in Figure 5E). Scale bar = 2 mm. **(D)** Representative images of heads and tongues measured from gross morphology (data shown in Figure 5E). Scale bar = 2 mm. **(E)** Measurement references for skeletal stains (data shown in Figure 5H-I and SF4F-H). **(F-H)** Quantification of skull features from skeletal staining of E18.5 fetuses that were unchanged across genotypes, with **(F)** skull length, **(G)** jaw length (multiple unpaired t-tests with Holm-Šidák correction, p = 0.048) and **(H)** cranial width. Error bars represent ± 1 SD. Abbreviations: snl = snout length, hl = head length, ctr = crown to rump, tl = tongue length, jl = jaw length, csa = cranial sagittal arc, sl = sagittal skull length, cw = cranial width.

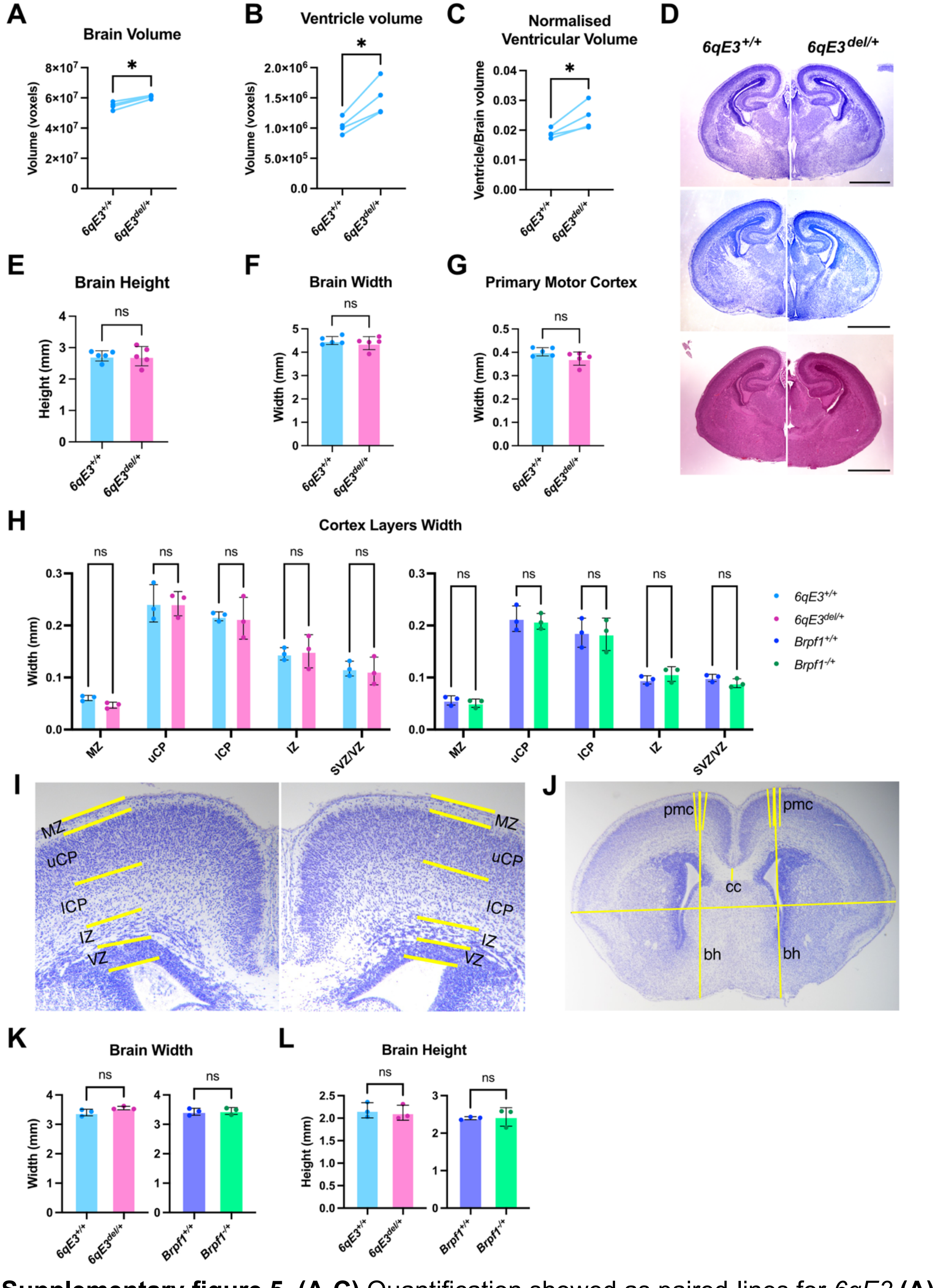


**Supplementary figure 5. (A-C)** Quantification showed as paired-lines for *6qE3* **(A)** brain volume, **(B)** ventricular volume and **(C)** ventricular volume normalised to brain volume, as displayed in Figure 6B-D. Error bars represent ± 1 SD. **(D)** *6qE3* E18.5 Nissl or H&E brain coronal sections of at approximately bregma 3.51 mm, with **(E-G)** showing measurements of **(E)** brain height (p = 0.9558), **(F)** brain width (p = 0.7282) and **(G)** primary motor cortex width (p = 0.2910). Multiple unpaired t-tests with Holm-Šidák correction, n = 5 per genotype. Error bars represent ± 1 SD. **(H)** Quantification of cortex layering for both mouse models at E18.5, at approximately bregma 2.19 mm, showing no changes in cortex layering. Multiple unpaired t-tests with Holm-Šidák correction, n = 3 per genotype, p > 0.05 for all. Error bars represent ± 1 SD. **(I)** Measurement references for cortex layering. (H-I) MZ = mesenchymal plate, uCP = upper cortical plate, lCP = lower cortical plate, IZ = intermediate zone, VZ = ventricular zone. **(J)** Measurement references for brain measurements shown in Figure 6F-G, and (K-L). Pmc = primary motor cortex, cc = corpus callosum, bh = brain height, bw = brain width. Note that where possible, measurements were taken for the left and right hemispheres, then averaged. For the primary motor cortex, 3 measurements were taken on each side and then averaged to the result displayed on the graph for each animal. **(K-L)** Quantification of **(K)** brain width and **(L)** brain height at approximately bregma 2.19 mm. Multiple unpaired t-tests with Holm-Šidák correction, n = 3 per genotype, p > 0.05 for all. Error bars represent ± 1 SD.

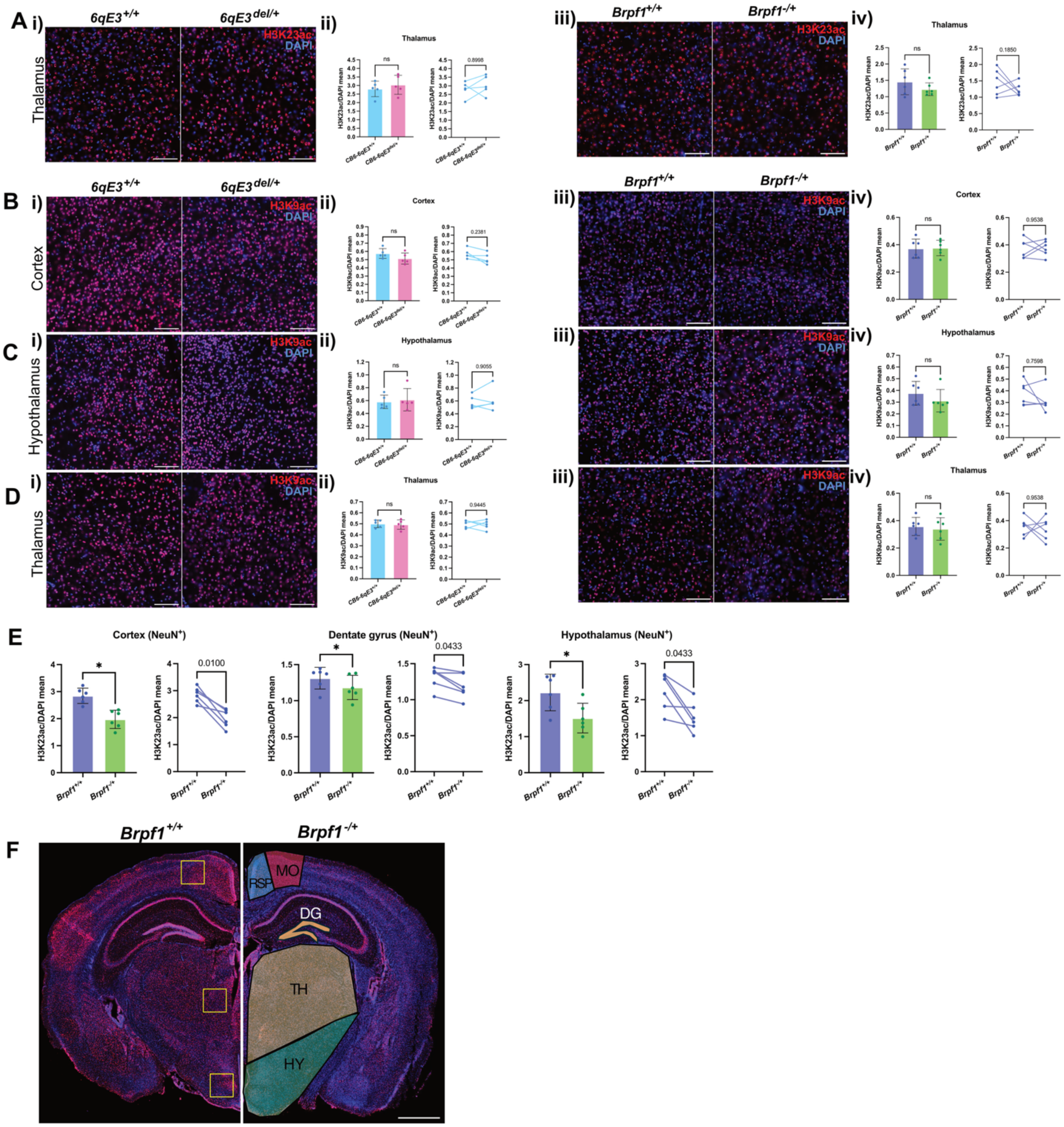


**Supplementary Figure 6.** Immunofluorescence of 16 um brain sections from 7-8 week old mice, with **(A)** showing H3K23ac in the thalamus. **(B-D)** H3K9ac in **(B)** the cortex, **(C)** the hypothalamus and **(D)** the thalamus. Scale bar = 100 µm. H3K23ac or H3K9ac appear in red, DAPI appears in blue. **i,iii.** Representative images of i. *Brpf1* and iii. *CB6-6qE3* samples. **ii,iv.** Quantification of acetylation levels in specific areas of the brain. Each dot represents an animal, with the mean intensity of the acetylation mark normalised to DAPI. Error bars represent ± 1 SD. Paired t-tests with Holm-Šidák multiple-testing correction. **(E)** H3K23ac levels for NeuN$^+$ cells in brain regions that showed significant reduction of acetylation. Error bars represent ± 1 SD. Paired t-tests with Holm-Šidák multiple-testing correction. **(F)** Measurement references for different areas of the brain (left), and labelled brain regions (right). Squares that were 550 um x 550 μm were placed in consistent regions across all samples, as indicated by yellow boxes. Scale bar = 1 mm.
